# Microstructural variation in the human striatum using non-negative matrix factorization

**DOI:** 10.1101/2021.06.10.447764

**Authors:** Corinne Robert, Raihaan Patel, Nadia Blostein, Chrisopher C. Steele, M. Mallar Chakravarty

**Author notes:** For correspondence, (FMS); (FS).

## Abstract

The striatum is a major subcortical connection hub that has been heavily implicated in a wide array of motor and cognitive functions. Here, we developed a normative multimodal, data-driven microstructural parcellation of the striatum using multiple magnetic resonance imaging-based metrics (mean diffusivity, fractional anisotropy, and the ratio between T1- and T2-weighted structural) from the Human Connectome Project Young Adult dataset (n=329 unrelated participants, age range: 22-35, F/M: 185/144). We further explored the biological and functional relationships of this parcellation by relating our findings to motor and cognitive performance in tasks known to involve the striatum as well as demographics. We identified 5 spatially distinct striatal components, for each hemisphere. We also show the gain in component stability when using multimodal versus unimodal metrics. Our findings suggest distinct microstructural patterns in the human striatum that are largely symmetric and that relate mostly to age and sex. Our work also highlights the putative functional relevance of these striatal components to different designations based on a Neurosynth meta-analysis.

## Introduction

The striatum is a deep grey matter nucleus known to be implicated in motor control (***Rolls, 1994***) and various executive and cognitive functions,including: goal-directed decision making (***Stott and Redish, 2014***; ***Haber et al., 2006a***), reward and motivation (***van den Bos et al., 2014***; ***Pauli et al., 2016***; ***Jung et al., 2014***; ***Haber et al., 2006a***), habitual motor learning (***Graybiel and Grafton, 2015***) and emotional regulation (***Hare et al., 2005***). Vari-ations in striatal structure and function have been implicated in various brain disorders including Parkinson’s disease (***Albin et al., 1989***; ***Hacker et al., 2012***), Huntington’s disease(***Rosenblatt and Leroi, 2000***), addiction (***Yager et al., 2015***; ***Li et al., 2015***; ***Graybiel and Grafton, 2015***), obsessive-compulsive disorders (***Graybiel and Rauch, 2000***; ***Shaw et al., 2015***; ***Milad and Rauch, 2012***), autism spectrum disorder (***Schuetze et al., 2016***), and schizophrenia (***Chakravarty et al., 2015***). Thus, the spatial subdivision of the striatum into regions informed by neuroanatomy is essential to relating striatal anatomy to function and behaviour. Previous parcellations of this important structure have leveraged magnetic resonance imaging (MRI) data using a combination of heuristic and contrast-based definitions (***Lehéricy et al., 2004***; ***Burrer et al., 2020***; ***Caravaggio et al., 2018***; ***Leh et al., 2007***). To overcome limitations inherent to these subjective definitions, data-driven parcellations based on structural connectivity (***Draganski et al., 2008***; ***Tziortzi et al., 2014***; ***Parkes et al., 2017***), resting-state functional connectivity, (***Jung et al., 2014***; ***Janssen et al., 2015***; ***Choi et al., 2012***; ***Marquand et al., 2017***)and task-based functional connectivity (***Pauli et al., 2016***) have been proposed. However, the existing parcellations have failed to characterize the tissue microstructure that necessarily constrains the organization and functional variation of the striatum.

In previous work, microstructural aspects of brain organization have been captured using structural and diffusion metrics derived from magnetic resonance imaging (MRI). Such microstructural metrics included the ratio between T1-weighted and T2-weighted images (T1w/T2w) (***Glasser and Van Essen, 2011***; ***Glasser et al., 2016***; ***Tullo et al., 2019***; ***Patel et al., 2020***; ***Tardif et al., 2016***), fractional anisotropy (FA) (***Alexander et al., 2007***; ***Lebel et al., 2008***; ***Patel et al., 2020***; ***Tardif et al., 2016***) and mean diffusivity (MD) (***Lebel et al., 2008***; ***Patel et al., 2020***; ***Tardif et al., 2016***). Typically these indices are used in isolation. The main goal of this study is to develop a data-driven microstructural parcellation of the striatum using a combination of T1w/T2w, FA and MD, and to link inter-individual variations in the obtained microstructural pattern to behavior and demographics. We will be using a framework previously developed and thoroughly investigated in ***Patel et al. (2020***) that used a similar approach to develop a multimodal parcellation of the human hippocampus using non-negative matrix factorization (NMF). The uncovered spatially distinct hippocampal parcels were found to be microstructurally distinct and stable across subjects.

We hypothesize that the decomposition of the covariance between the T1w/T2w, FA and MD metrics should yield parcels of the striatum that are more stable across our subjects relative to a decomposition based on a single metric. (***Sotiras et al., 2015***; ***Patel et al., 2020***). Another goal of this study is to relate inter-individual variations in the obtained microstructural parcels to motor and cognitive performance. Finally, we aim to relate group-level microstructural patterns of the striatum to brain function through a functional MRI (fMRI) based platform called Neurosynth (***Yarkoni et al., 2011***).

## Methods and Materials

### Overview

A schematic illustration of the methods of analyses used in the present study can be found in ***Figure 1***. We used structural and diffusion MRI data from the Human Connectome Project (Data). The striatum segmentations were generated automatically using the Multiple Automatically Generated Templates (MAGeT) Brain algorithm (Automatic striatum segmentation). A population average constructed using the T1w and T2w images of each subjects was also generated to provide a common space for the microstructural metrics used in our analyses (Population average). The obtained striatal labels, T1w/T2w, FA and MD maps to this common space to construct the input matrices that then under-went NMF decomposition (Implementation, ***Figure 1*A. B**, &C). A stability analysis was performed to find the optimal number of components (Stability analysis) and the final solution was compared to unimodal solutions. The final multi-modal NMF solution was used to generate neuroanatomically distinct clusters that are used to describe the microstructural anatomy of the striatum (Non-negative matrix factorization, ***Figure 1*D**). Then, we use the inter-indiviudal variability in the striatal components, characterized by the NMF subject-level weights (Non-negative matrix factorization, ***Figure 1*E**) to understand how patterns of covariance may relate to behaviour and demographics using Partial Least Squares correlation analysis (Microstructure-behaviour relationships, ***Figure 1*G**).Finally, to ascertain their putative functional relevance, these clusters were used as input to Neuro-synth meta-analytical decoder to compare them to meta-analyzed fMRI findings (Neurosynth image decoder, ***Figure 1*F**).

**Figure 1.**
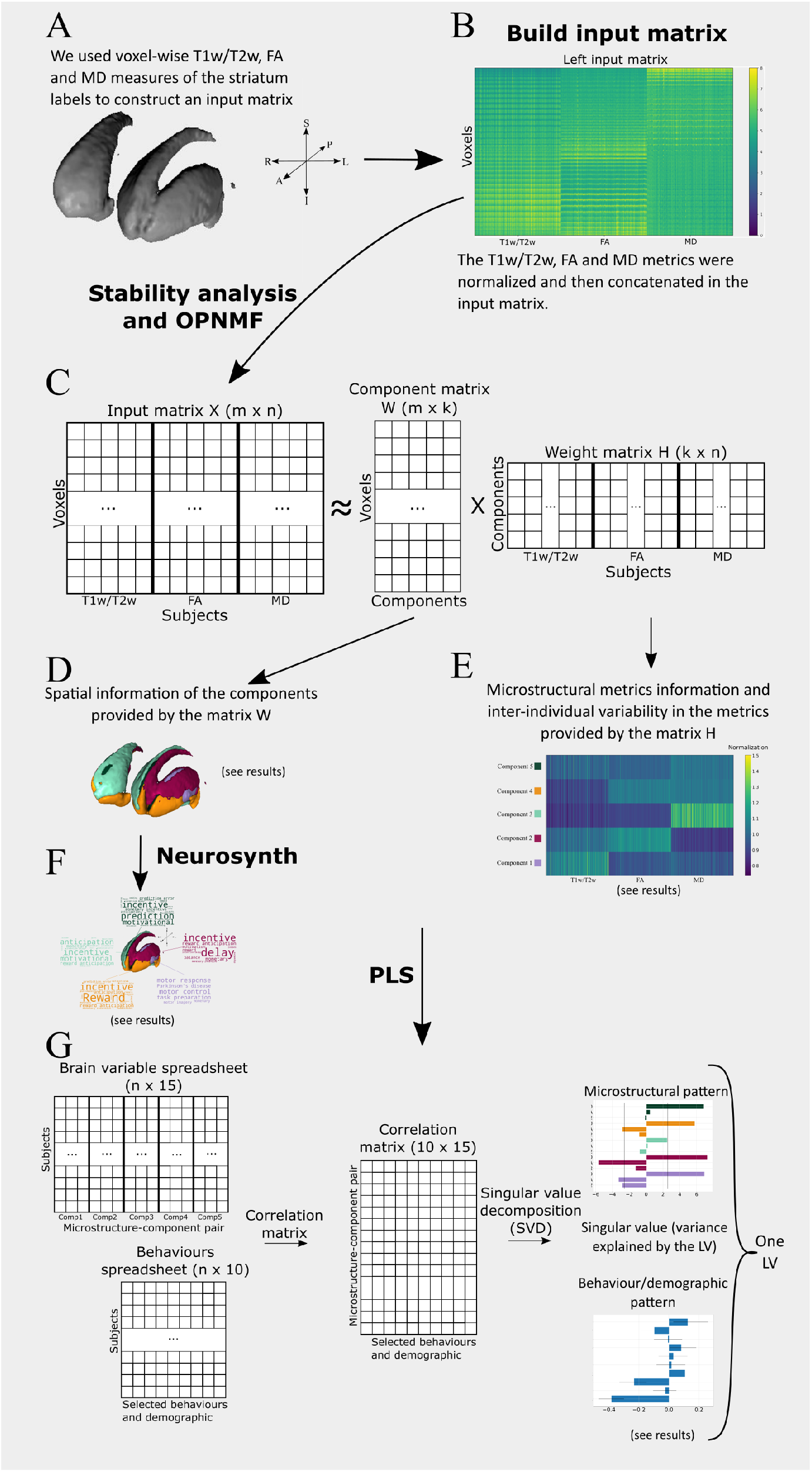
A) We used the chosen microstructural metrics in automatically segmentated striatum labels (obtained with the MAGeT Brain algorithm) of our subjects in the a constructed common space B) We concatenated the striatal voxels in column vectors of all our subjects to build an input matrix. The left and right input matrix were build independently. C) We extracted spatially distinct components representing patterns of covariance in microstructure across subjects using orthogonal projective non-negative matrix factorization (OPNMF). OPNMF decomposes an input matrix into a component matrix W and a weight matrix H. As OPNMF extracts a predefined number of component k, we performed a stability analysis to assess the accuracy and spatial stability at each granularity from 2 to 10 (see ***Figure 2*A**). D) The component matrix W describes how much each voxel weight into a specific component providing spatial information about the clusters. F) We related each component to functional MRI findings by using theNeurosynth reverse-inference framework that meta-analytically relates striatal components to psychological states. E) The weight matrix H contains the weight of each subject’s metrics onto each component, describing microstructural variation in the metrics found in the input matrix (T1w/T2w, FA, MD) between subjects. G) We used Partial Least Squares (PLS) analysis to identify patterns of covariance between the striatal components T1w/T2w, FA and MD proportions with behavioural and demographic data. PLS is a multivariate technique that analyses the association between our component-metric pairs (leftmost top) and selected behaviour/demographics (leftmost bottom) variables resulting in a set of LVs. The significance of the covariance patterns uncovered by the LVs was assessed using permutation testing while the reliability of each brain specific weight was assessed using bootstrap sampling.

### Data

We used multimodal MRI along with behavioural and demographic data from the Human Connectome Project (HCP) Young Adult dataset. We selected structural and diffusion MRI data from 333 unrelated subjects (from a cohort of 1086 twin and non-twin siblings) with age ranging from 22-35 years (***Van Essen et al., 2013***). Most of the participants were individuals born in Minnesota and participants were excluded for severe neurodevelopmental, neuropsychiatric or neurologic disorders (***Van Essen et al., 2013***). All structural and diffusion MRI data were acquired on a customized Siemens 3T Skyra scanner with a 100 mT/m gradient (***Van Essen et al., 2013***).

#### T1w/T2w images

We used preprocessed T1 (T1w)- and T2-weighted (T2w) images from the [HCP database](https://db.humanconnectome.org/) (0.7 mm isotropic images) (***Van Essen et al., 2013***). T1w images were further preprocessed using the minc-bpipe library minc-bpipe library to perform intensity non-uniformity correction, cropping of the neck region and brain mask generation. T1w images were used to derive a minimally-biased group template (as described below) and the T1w/T2w images were used as a putative measure of voxel-wise myelin content (***Glasser and Van Essen, 2011***; ***Tullo et al., 2019***). Detailed preprocessing of the HCP data is described in detail elsewhere (***Van Essen et al., 2013***; ***Glasser et al., 2013***).

#### DWI scalars

The preprocessed diffusion weighted imaging data (1.25 mm isotropic voxel dimensions) were also downloaded via the HCP online portal. The processing pipeline applied to the diffusion data by the HCP is described in ***Glasser et al.*** (***2013***). The diffusion data were further processed by R.P. in another study from our group (***Patel et al., 2020***) with [MRtrix](https://github.com/MRtrix3/mrtrix3) (***Tournier et al., 2012***) to estimate MD and FA maps for each subject. To do so, single shell 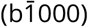 data was used to construct the tensor with weighted least-squares (***Basser et al., 1994a***) and iterated least-squares (***Veraart et al., 2013***) using the dwi2tensor command. Then, the MD and FA maps were estimated from the tensor using the tensor2metric command (***Basser et al., 1994b***; ***Westin et al., 1997***).

#### Automatic striatum segmentation

The striatum was segmented in each subject’s T1w image using the publicly available [MAGeT brain algorithm](https://github.com/CobraLab/MAGeTbrain) (***Chakravarty et al., 2013***). We used [5 high-resolution manually segmented subcortical atlases](https://github.com/CoBrALab/atlase) based on the reconstruction of serial histological data (***Chakravarty et al., 2006***; ***Tullo et al., 2018***). All registrations in this section and the next section were performed using the [Automatic Normalization Registration Tools (ANTs)](https://github.com/vfonov/mincANTS) (***Avants et al., 2010***). Two runs of MAGeT brain were performed by N. B. on the entire HCP cohort (N=1086): manual quality control of the outputs from the first run allowed for the selection of the 21 subjects with the best segmentations; these subjects were then used as templates for the second and final run. This allowed for more subjects to pass manual quality control for output quality (see the [guide](https://github.com/CobraLab/documentation/wiki/MAGeT-Brain-Quality-Control-(QC)-Guide)). MAGeT brain was run separately for each hemisphere to account for anatomical asymmetries and to improve segmentation accuracy.

#### Population average

A population average was used to obtain a voxel-wise correspondence between our 333 subjects and was computed by R.P. in another study from our group (***Patel et al., 2020***). We used the transformation files from the T1w images to common space to warp each subject’s striatum segmentation, T1w/T2w, FA and MD images to the common space using the antsApplyTransforms command. T1w/T2w images were filtered using a Gaussian weighted average to remove any outlier values (***Patel et al., 2020***; ***Glasser and Van Essen, 2011***).

Striatum labels that passed quality control (left, n=252; right, n=289) were transformed to the common space and a unified label was generated by voxel-wise majority vote. The final labels were adjusted for over-segmentation in areas such as the lateral ventricle or the internal capsule (see examples [here](https://github.com/CobraLab/documentation/wiki/MAGeT-Brain-Quality-Control-(QC)-Guide)) to minimize partial voluming effects of ventricles.

### Non-negative matrix factorization

We used an orthonormal projective variant of non-negative matrix factorization (OPNMF). This method provides a part-based decomposition of the input variables while prioritizing sparsity in the solution (***Yang and Oja, 2010***; ***Sotiras et al., 2015***). OPNMF has already been proven effective in estimating covariance patterns in neuroimaging data while providing an easier interpretation of the results than other matrix decomposition techniques such as principal component analysis (PCA) or independent component analysis (ICA) (***Sotiras et al., 2015***). Briefly, NMF decomposes an input matrix (m × n) into two matrices; a component matrix *W* (m × k) and a weight matrix *H* (k × n) where k is the number of components that needs to be specified by the user, m is the number of striatal voxels and n is the number of subjects (329) for the unimodal implementation and the number of subject-metric pairs (329*3) for the multimodal implementation. Here we use the same nomenclature as in ***Patel et al.*** (***2020***). As we are using the orthogonal projective version of NMF (OPNMF), our decomposition identifies k spatially distinct patterns of covariance across voxels (found in W) and across subjects and metrics (found in H). We describe below how we implemented OPNMF as well as how we interpreted the decomposition results. More theoretical concepts about OPNMF and its implementation can be found in the supplements. We examined each microstructural measure (T1w/T2w, FA and MD) separately through a unimodal implementation of OPNMF and simultaneously through a multimodal implementation of OPNMF. More details on the implementation of the unimodal and multimodal OPNMF analyses are described below.

### Implementation

#### Input matrices

We used the fused left and right average striatum labels (Population average) to perform a ROI-based extraction for the T1w/T2w, FA and MD metrics using the [TractREC package](https://github.com/CoBrALab/TractREC). For each subject, the voxels of the striatum labels were extracted and stacked into a column vector of size (# striatal voxels × 1 subject). Therefore, we obtained voxel-wise column vectors for each subject and each of the microstructural metric (T1w/T2w, FA and MD). Hence, we obtained 3 metric vectors per hemisphere for every subject, resulting in 6 column vectors per subject. As OPNMF was applied on the two hemispheres separately, the left and right input matrices for the unimodal and multimodal OPNMF were constructed independently.

For the unimodal input matrices, we concatenated the 329 corresponding column vectors to obtain 6 (# striatal voxels × 329 subjects) matrices (per hemisphere and metric). The unimodal matrices were normalized using a standard z-score and shifted by the minimum value to obtain non-negativity.

For the multimodal matrices, we concatenated the unimodal matrices that were normalized to account for different scales of magnitude, resulting in one (striatal voxels × 3 × 329) matrix per hemisphere. We then shifted all the values in our multimodal input matrices by the minimum value.

Once the input matrices were constructed, we applied the OPNMF algorithm on the left and right striatum separately. We used MatLab R2016a and some [OPNMF matlab functions](https://github.com/asotiras/brainparts) (***Sotiras et al., 2015***; ***Boutsidis and Gallopoulos, 2008***; ***Halko et al., 2011***; ***Yang and Oja, 2010***). The OPNMF algorithm was initialize using non-negative double singular value decomposition (SVD) and the following hyperparameters: max iterations = 100000 and tolerance = 0.00001 as in (***Patel et al., 2020***).

#### Interpretability

OPNMF outputs a component matrix *W* and a weight matrix *H*. The (striatal voxels × component matrix *W* describes how much each voxel contribute to a specific component. The (k × (3 × 329 subjects)) weight matrix *H* presents the loading of each subject’s metrics onto each component, describing microstructural variation in T1w/T2w, FA and MD between subjects.

The properties of OPNMF enable us to cluster voxels via a winner take all approach of each voxels component scores, such that each voxel was assigned to a single cluster for which it had the highest component score. Therefore, *W* provides spatial information about the striatal components.

For the weight matrix *H*, we have the subject-metric pairs as columns, component as rows and every entry represents the proportion of the metric-subject pair that contributes to each component. For a given component, the weight of a microstructural metric should be similar across subjects with some variability. Hence *H* conveys information about how much T1w/T2w, FA and MD contribute to each striatal component as well as how those proportions vary across individuals within each component.

### Stability analysis

To select the optimal number of components, a stability analysis was run to assess the accuracy and spatial stability at each granularity from 2 to 10 (***Patel et al., 2020***). The stability analyses for the left and right striatum were performed independently.

We split our 329 subjects into two groups (a and b) of size n*_a_* = 164 and n*_b_* = 165 that were stratified by age. We repeated this procedure to create 10 different splits, to obtain 10 × 2 = 20 groups. For each split we created multimodal input matrices *X_a_* and *X_b_* as described in Implementation and ran OPNMF on each split independently (resulting in 20 × 2 × 9 = 360 runs). For each split and granularity we obtained two-component matrices *W_a_* and *W_b_* which are of dimension (#striatal voxels × granularity) and two weight matrices *H_a_* and *H_b_* of dimensions (granularity × (n*_a_* or n*_b_*)). The reconstruction error for each split was computed as follows:

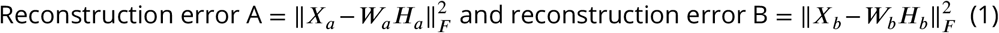

Where *X_a_* and *X_b_* are the input matrices of two respective groups in a split. We reported the gradient reconstruction error that corresponds to the change in the reconstruction error from a granularity *k* to the granularity *k* + 1. Hence, the gradient reconstruction error was computed by subtracting reconstruction error matrix of the granularity *k* + 1 with the reconstruction error matrix of the granularity *k* and then averaging all the differences. Then we average over all splits to get reconstruction error average and standard deviation for every granularity.

The accuracy is computed for each split by first taking two similarities matrix *CW_a_* and *CW_b_* of dimensions (# striatal voxels × # striatal voxels). *CW_ij_* contains the cosine similarity between the components scores of voxel *i* and voxel *j*. If cosine similarity is high, it means that voxels *i* and *j* have similar component scores and that they are likely in the same cluster (***Patel et al., 2020***). Hence, a row *i* in the matrix *CW_a_* represents the similarity of the voxel *i* with all the other voxels for the group a. This is the same for the matrix *CW_b_*. Then, we computed the correlation between corresponding rows of *CW_a_* and *CW_b_*, to know if a certain voxel *i* is similar to the same group of voxels when OPNMF is applied on another group (***Patel et al., 2020***). If the correlation between corresponding rows of voxels was high, we conclude that the stability was high for this voxel (stability coefficient close to 1). On the other hand, instability (stability coefficient close to −1) was implied by a low correlation between corresponding rows of voxels. Finally, we took the average for all voxels and we repeated this procedure for each split to get the average and standard deviation stability coefficient for every granularity.

To assess the benefit of using multimodal data versus unimodal data, we carried out a unimodal stability analysis for the T1w/T2w, FA and MD metrics separately. As for the multimodal stability analyses, the left and right unimodal stability analyses were conducted separately, for a total of 6 unimodal stability analyses.

### Microstructure-behaviour relationships

To link inter-individual variation in striatal OPNMF components to behaviour and demographics, we sought to examine their relationship to a set of behaviours and demograph-ics available from the HCP by using subject-level weights as a measure of their specific microstructural loadings (in matrix *H* for each component, see Implementation). We considered all the motor-related behaviours available in the HCP test battery, as the relationship between the striatum and motor function is well known (***Mink, 1996***; ***Delong et al., 1983***). This included endurance (NIH Toolbox 2-minute Walk Endurance Test), locomotion (NIH Toolbox 4-Meter Walk Gait Speed Test), dexterity (NIH Toolbox 9-hole Pegboard Dexterity Test) and strength (NIH Toolbox Grip Strength Test). We also considered cognitive tests related to impulsivity (***Hariri et al., 2006***; ***Buckholtz et al., 2010***; ***Dalley et al., 2008***), motor inhibition and cognitive control (***Vink et al., 2005***; ***Schouppe et al., 2014***). Impulsivity was assessed using the delay-discount task (DD) (***Green et al., 2007***; ***Estle et al., 2006***) with the area under the curve (AUC) of DD as a summary measure. Low values for the AUC suggests delayed rewards are less valuable to the subject and vice versa (***Myerson et al., 2001***). Motor inhibition and cognitive control were measured by the HCP using the Flanker task from the NIH toolbox (***Schouppe et al., 2014***). We also considered age in years, years of education and gender as demographic measures.

### Partial Least Squares

To associate the selected behaviours to the subjects’ metric-wise component weightings, we used Partial Least Squares Correlation ([PLSC](http://pls.rotman-baycrest.on.ca/UserGuide.htm)). PLSC is a multivariate statistical technique that analyses the association between two sets of high-dimensional variables(***Krishnan et al., 2011***; ***Zeighami et al., 2019***; ***McIntosh and Lobaugh, 2004***; ***Patel et al., 2020***). In the context of the current study, we related the set of individual component weightings obtained from the H matrix in OPNMF (brain data) to the set of behavioural/demographics variables mentioned above (behaviour data).

Briefly, in PLSC, major patterns of covariance are extracted from the correlation matrix of our to initial sets using SVD (***Krishnan et al., 2011***; ***Zeighami et al., 2019***). The SVD decomposition yields a set of uncorrelated latent variables (LVs) (***Krishnan et al., 2011***; ***Zeighami et al., 2019***). Each LV has a singular value, which is the proportion of covariance explained by this LV. There are also a set of brain scores and behavioural scores, describing the extent to which brain and behavioural elements are contributing to the LV on a per-subject basis (***Krishnan et al., 2011***; ***Zeighami et al., 2019***).

We assessed significance of each LV using non-parametric permutation testing on the singular values. The stability of the individual brain and behavioural scores elements or weight were assessed by using bootstrap sampling (***Krishnan et al., 2011***; ***Zeighami et al., 2019***; ***McIntosh and Lobaugh, 2004***; ***Patel et al., 2020***).

#### Implementation

Here, the brain matrix had dimensions (329 subjects × 3 metrics × k components) with one row for each subject and one column for each component-metric pair. The behavioural variables were stored in a 329 × 10 matrix, with the subjects as rows and the performance of selected behavioural tests along with age, sex (coded as 0/1 for M/F) and years of education as columns. Our PLSC outputs represent a pattern of covariance between the selected behaviours and component-wise microstructural data. For the permutation testing, we computed 10000 permuted brain matrices to construct a null distribution of singular values. We considered a threshold of P<0.05 to be significant, as it corresponds to a 95% confidence that the singular value of the original LV is higher than the singular value of the permuted LV (***Patel et al., 2020***). As for the bootstrap sampling, we generated 1000 bootstrap samples and considered a brain salience weight with BSR*>*2.58 to be significant as it corresponds to P<0.01 (99% confidence) (***Krishnan et al., 2011***; ***McIntosh and Lobaugh, 2004***; ***Patel et al., 2020***).

### Neurosynth image decoder

We related each component to functional MRI findings by using the Neurosynth association test framework that meta-analytically relates striatal components to brain function (***Yarkoni et al.*** (***2011***)). The [Neurosynth database](https://neurosynth.org/) is comprised of meta-analytic functional maps for 1335 terms automatically generated from 14371 studies. Through the Neurosynth [Image Decoder](https://neurosynth.org/decode/), it is possible to compare any brain map to the entire Neurosynth database and thus quantitatively infer cognitive states for each uploaded map (***Yarkoni et al., 2011***; ***Chang et al., 2012***). More specifically, provides posterior probability maps associated with a given term representing the likelihood that this term is being used in a study if activation is observed in the striatal voxels that we provided (see [association test](https://neurosynth.org/faq/)). As our striatal components were in the previously computed common space (Population average), we warped the components to MNI space before uploading them one by one to NeuroVault as ROI-based NIFTI images. Our [MNI space striatal components](https://identifiers.org/neurovault.collection:9498) are publicly and can be used for further analysis. From the posterior probability maps provided by Neurosynth, we excluded maps with anatomical keywords to focus on cognition related terms. We also excluded maps with keywords that were either unspecific, such as “life”, or redundant like “loss” and “losses”.

## Results

### Data

The final sample size included 329 subjects from the Human Connectome Project Young Adult dataset. The demographic information of our participants is displayed in ***Table 1***. We note that there is a significant difference in the mean age between males and females (*t*(327) = 3.1, p<0.05), and there is no significant difference between males and females in handedness (*t*(327) = 1.4, p>0.1) and overall cognition (*t*(327) = 1.4, p*>*0.1).

**Table 1.** Participants demographics. MMSE: score on 30 of the Mini-mental state examination

| Sex | Number | Mean age (years) | Mean Handedness | Mean overall cognition (MMSE) |
| --- | --- | --- | --- | --- |
| Females | 185 | 29.01±3.63 | 66.59±47.31 | 29.18±0.97 |
| Males | 144 | 27.71±3.68 | 59.69±44.20 | 29.01±1.08 |
| Overall | 329 | 63.57±46.06 | 63.57±46.04 | 29.11±1.02 |

### Stability analysis

The results of the stability analysis are shown in ***Figure 2***. In ***Figure 2*A**, the stability coefficient (red) of the multimodal OPNMF decomposition is displayed for the left and right striatum, as well as the gradient of the reconstruction error (blue) for all chosen granularities. In the right striatum, there is a net drop in the stability of the OPNMF clusters at *k* = 4, while the stability of the left OPNMF clusters slightly decay for *k* ≥ 3.

**Figure 2.**
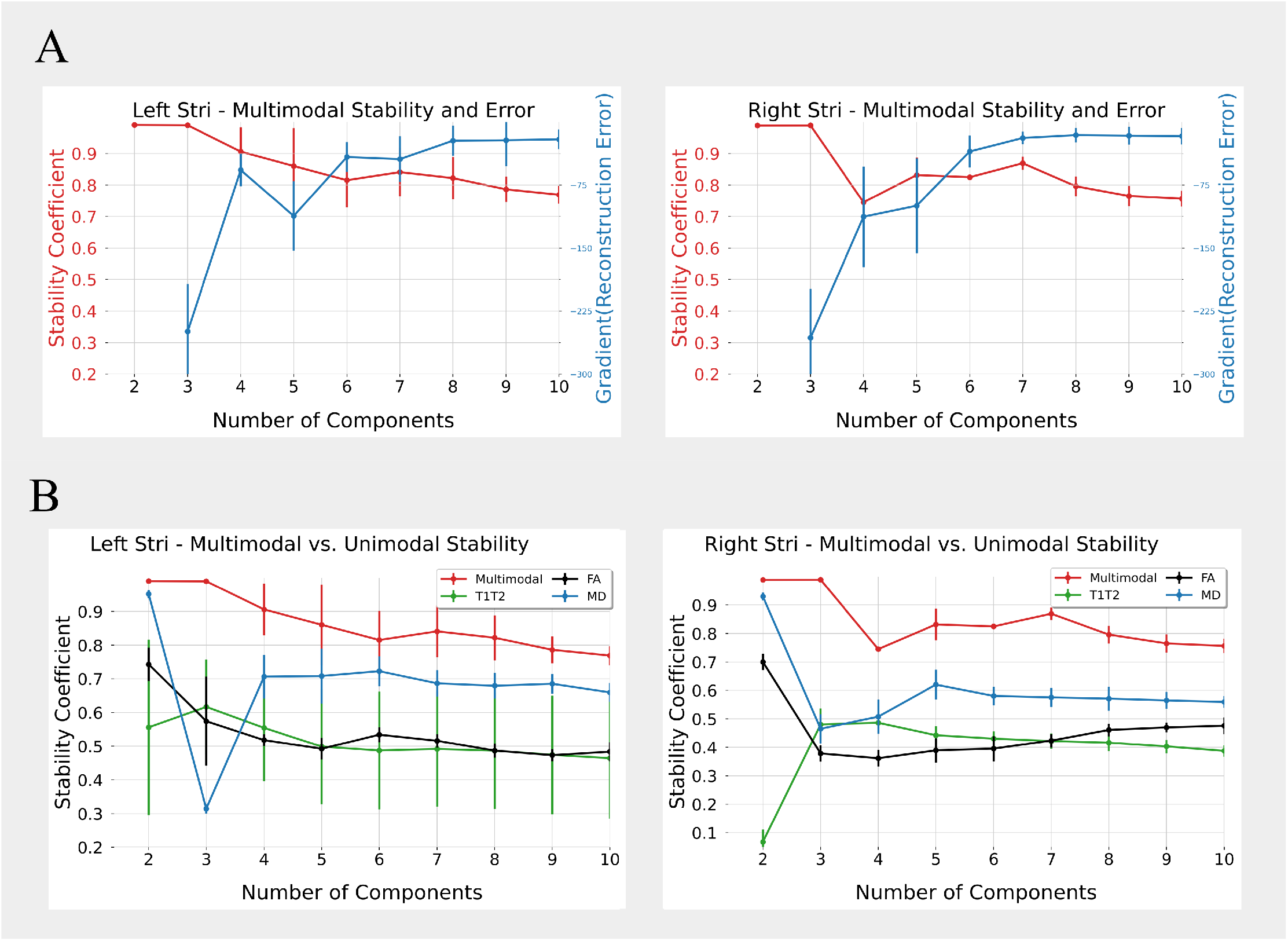
A) Stability score and gradient reconstruction error when performing NMF using 2 to 10 clusters. As we want to maximize the stability while minimizing the reconstruction error, we chose to use 5 components for the rest of the analysis. B) Comparison of the stability score of NMF on multimodal data (a combination of T1w/T2w, FA and MD (red)) versus unimodal data (either only T1w/T2w (green), only FA (black) or only MD (blue)) using 2 to 10 clusters.

The gradient reconstruction error increases as the granularity increases for both hemispheres. The gradient reconstruction error going from *k* = 3 to *k* = 4 increases dramatically for both the left and right striatum, suggesting that there is more gain from going to *k* = 2 to *k* = 3 components than from *k* = 3 to *k* = 4 components. However, the gain in the reconstruction error of the left striatum is better than expected when going from *k* = 4 to *k* = 5. The plateau in the reconstruction error for *k*6 in both hemispheres suggests that major patterns of covariance have been captured. Hence, *k* = 5 was chosen as the optimal number of components for the left and right striatum as it is the granularity that provides the best balance between the stability coefficient and the reconstruction error (accuracy) of the OPNMF multimodal decomposition.

The results of the stability analysis comparing the multimodal versus the unimodal OPNMF decomposition with k ranging from 2 to 10 is shown in ***Figure 2*B**. The stability coeicient of the unimodal metrics T1w/T2w (green), FA (black) and an MD (blue) is lower than the stability achieved with the multimodal decomposition (red) for both hemispheres. Due to the gain in stability of the multimodal decomposition, we decided to only conserve the 5-component multimodal solution for further analysis.

### Striatal components

***Figure 3*A** shows a 3D representation of the left and right striatal components, while ***Figure 3B*** displays selected labelled and unlabelled coronal slices. The weight matrix in ***Figure 3*C** shows the metrics proportion in each component. We only show the left weight matrix as it is almost identical to the right weight matrix. The weight matrix was divided by the mean within rows to offer better visualization of within component variation in the microstructural metrics.

- Component 1 (lilac in ***Figure 3*A**&B) is characterized by higher values of T1w/T2w compared to MD and FA with slighty lower values of FA compared to the previous metrics (first row from the bottom in ***Figure 3*C**). Component 1 includes the dorsal putamen as well as the dorsolateral caudate nucleus.
- Component 2 (dark magenta in ***Figure 3*A**&B) is characterized by a high proportion of FA, followed by T1w/T2w and MD (second row from the bottom in ***Figure 3*C**). Component 2 forms a thin capsule around the dorsal putamen and also includes the exterior lateral caudate next to the internal capsule.
- Component 3 (light mint in ***Figure 3*A**&B) is characterized by high MD metrics compared to the proportion of T1w/T2w and FA (third row from the bottom in ***Figure 3*C**). Component 3 is a thin cluster including the anterior and posterior medial caudate nucleus along the anterior horn of the lateral ventricle.
- Component 4 (orange in ***Figure 3*A**&B) is characterized by lower T1w/T2w values compared to FA and MD (fourth row from the bottom in ***Figure 3*C**). Both FA and MD in component 4 are slightly above average. This component includes the nucleus accumbens and a part of the outer ventrolateral putamen.
- Component 5 (dark green in ***Figure 3*A**&B) is characterized by lower values of FA compared to the values of T1w/T2w and MD in this component (last row from the bottom in ***Figure 3*C**). Component 5 includes the inner anterior ventral caudate, the medial caudate body and some part of the ventral putamen.

**Figure 3.**
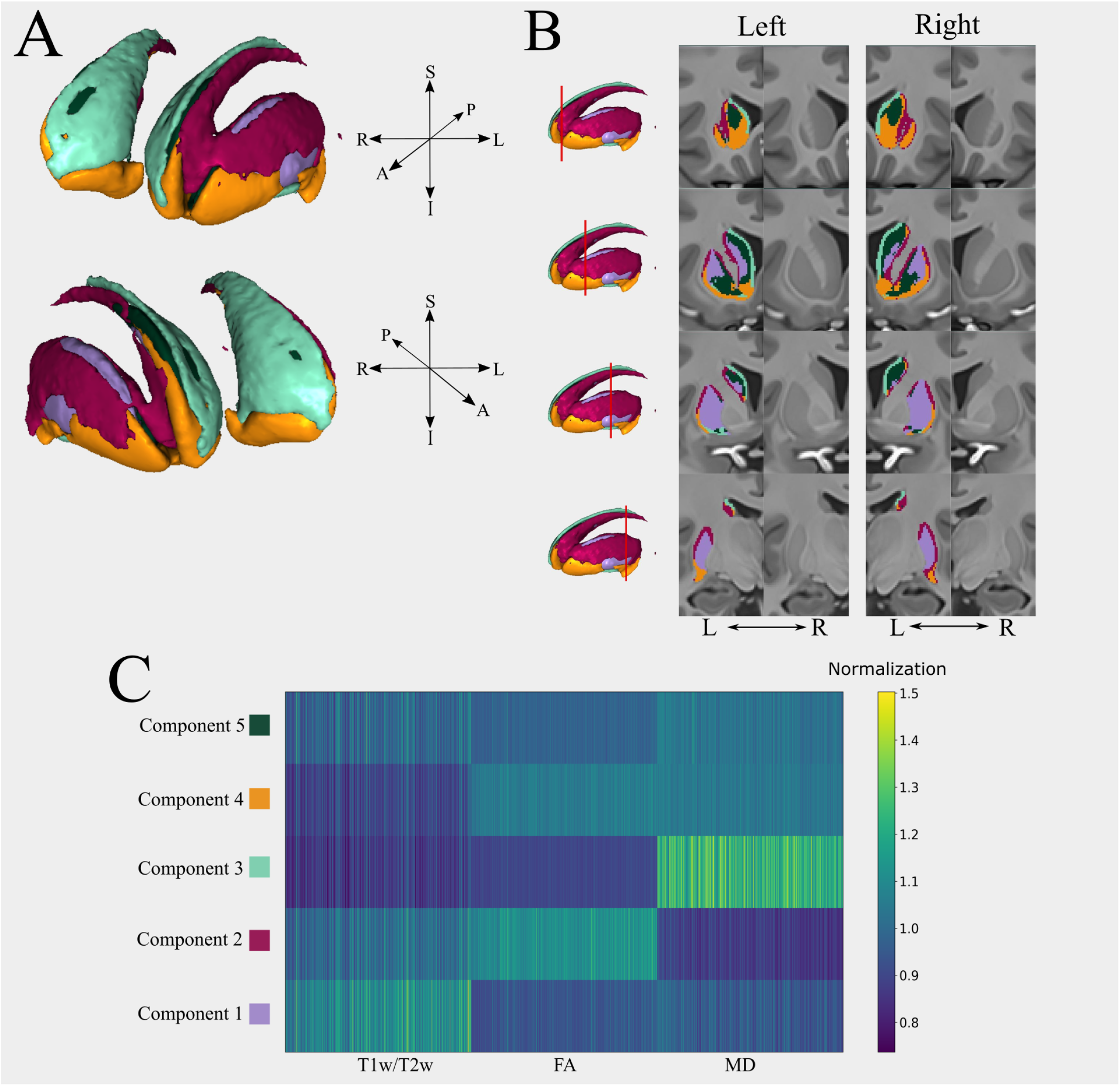
A) 3D rendering of the 5 components solution (A: anterior, P: posterior, S: superior, I: inferior, R: right, L: left). B) Coronal slices showing the labelled and unlabelled (side-by-side columns) left and right striatum. C) Weight matrix output from NMF of the left striatum, showing how the microstructural metrics weight into each component (the right weight matrix is almost identical). For the normalization, we divided each component (row in the matrix) by the mean value in that specific component to show within component variation in the microstructural metrics.

### Partial Least Square analysis

To relate individuals subject’s weighting from the weight matrix of OPNMF to selected behaviours and demographics, we used Partial Least Square correlation analysis on the left and right hemisphere independently. Using permutation testing, we identified four significant latent variables, two for the left striatum (p<0.05) and two for the right striatum (p<0.05) shown in ***Figure 4***. ***Figure 4*A** shows the behavioural patterns associated with the LV where the y-axis shows the behaviour and demographic measures and the x-axis shows the correlation of that behaviour/demographic and the LV. ***Figure 4*B** shows the microstructual patterns associated with the LV where the y-axis shows the componentmetric pairs and the x-axis denotes the bootstrap ratio (BSR).

**Figure 4.**
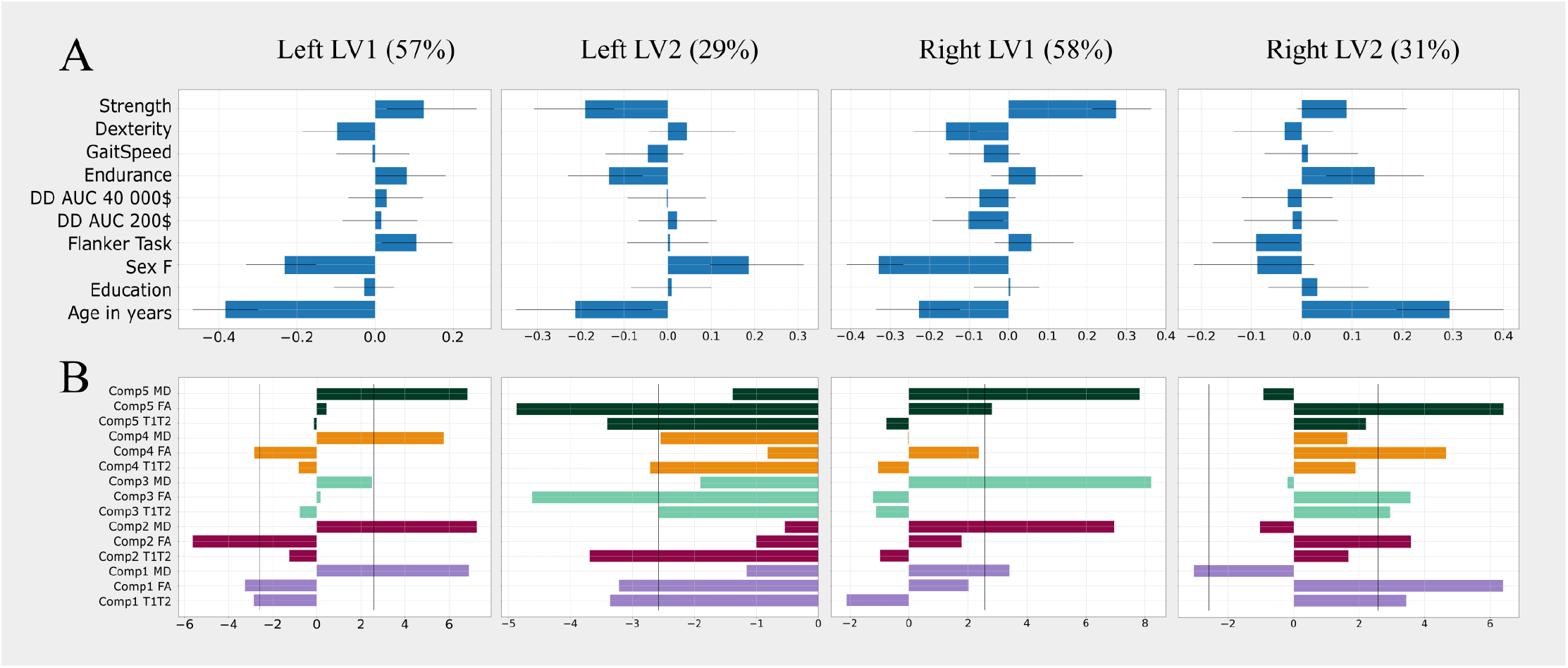
Results of the PLS analysis, we show only the latent variables (LVs) that were significant (p<0.05). The percentage next to the LV’s name corresponds to the covariance explained by this LV. A) Behavioural patterns of the left LV1 (first column), left LV2 (second column), right LV1 (third column) and right LV2 (fourth column). The y-axis denotes the behavioural and demographics measures used in the analysis (DD AUC: Delay discounting area under the curve), while the x-axis corresponds to the correlation of the behaviours with the LV. B) Microstructural patterns associated with the four significant LVs identified. Here,the y-axis correspond to the component-metric pairs and the x-axis denotes the bootstrap ratio (BSR). The black line in the microstructural patterns graph represent a BSR of 2.58 (equivalent to a 99% C.I.). The colors of the bars are associated with the component (see ***Figure 3***C).

The first left LV (left LV1; ***Figure 4*A** top row) explains 57% of the covariance between our two and was associated with young age (R=−0.383, 95% C.I.=[−0.467,−0.300]), male sex (R=−0.232, 95% C.I.=[−0.330,−0.150]), increased average performance on the Flanker task (R=0.107, 95% C.I.=[0.018,0.199]), increased strength (R=0.125,95% C.I.=[0.030,0.261]) and decreased dexterity (R=−0.097,95% C.I.=[−0.185,−0.013]). The correlated microstructural features include increased MD across all 5 components, decreased FA in components 1,2 and 4 and decreased T1w/T2w in component1.

Left LV2 (***Figure 4*A** bottom row) explains 29% of the variance and is associated with lower age (R=−0.213, 95% C.I.=[−0.350,−0.035]), female sex (R=0.186, 95% C.I.=[0.097,0.314]), decreased strength (R=−0.190, 95% C.I.=[−0.308,−0.124]) and endurance (R=−0.136, 95% C.I.=[−0.230,−0.058]). The correlated microstructural features include decreased FA in components 1, 3, 5 and decreased T1w/T2w across all components.

Right LV1 (***Figure 4*B** top row) explains 58% of the variance and was mainly driven by younger age (R=−0.227, 95% C.I.=[−0.335,−0.122]), male sex (R=−0.329, 95% C.I.=[−0.410,−0.267]), increased strength (R=0.2737,95% C.I.=[0.213,0.362]), decreased dexterity (R=− 0.1571, 95% C.I.=[−0.241,−0.079]) and AUC for both delay discounting measures DD AUC 200$(R=−0.1015, 95% C.I.=[−0.191,−0.013]) and DD AUC 40 000$(R=−0.0734,95% C.I.=[−0.160, 0.019]). The correlated microstructural features include increased MD across all components.

Right LV2 (***Figure 4*B** bottom row) explained 31% of the variance and was associated with young age (R=0.293, 95% C.I.=[0.188,0.400]), male sex (R=−0.09, 95% C.I.=[−0.213,0.025]), increased endurance (R=0.145, 95% C.I.=[0.049,0.243]) and below average performance in the Flanker task (R=−0.090, 95% C.I.=[−0.177,−0.004]). The correlated microstructural features included increased FA across all components, increased T1w/T2w in components 1 and 3 and decreased MD in component 1.

### Decoding with Neurosynth

The results of the association test performed by Neurosynth for the left and right striatal components are in ***Figure 5***. Some posterior probability maps were unique for certain components. Posterior probability maps with keywords related to motor function such as “motor control” and “motor response” were only associated with the first component of the left and right striatum. The same result was found for maps related to Parkinson’s disease. The posterior probability map associated with the keyword “age” was only associated with the fifth right striatal component. In general, the correlations obtained for right striatal components were smaller than the correlations obtained for the left striatal components. There was also a lot of overlap within the set of posterior probability maps across components and hemispheres, but the correlation values associated with similar maps was different between components. For instance, in ***Figure 5*** top, we can see that both the left component 3 (light green) and the left component 4 (orange) are associated both associated with the “reward” posterior probability map. However, the left component 4 has a bigger correlation with the “reward” map then the left component 3.

**Figure 5.**
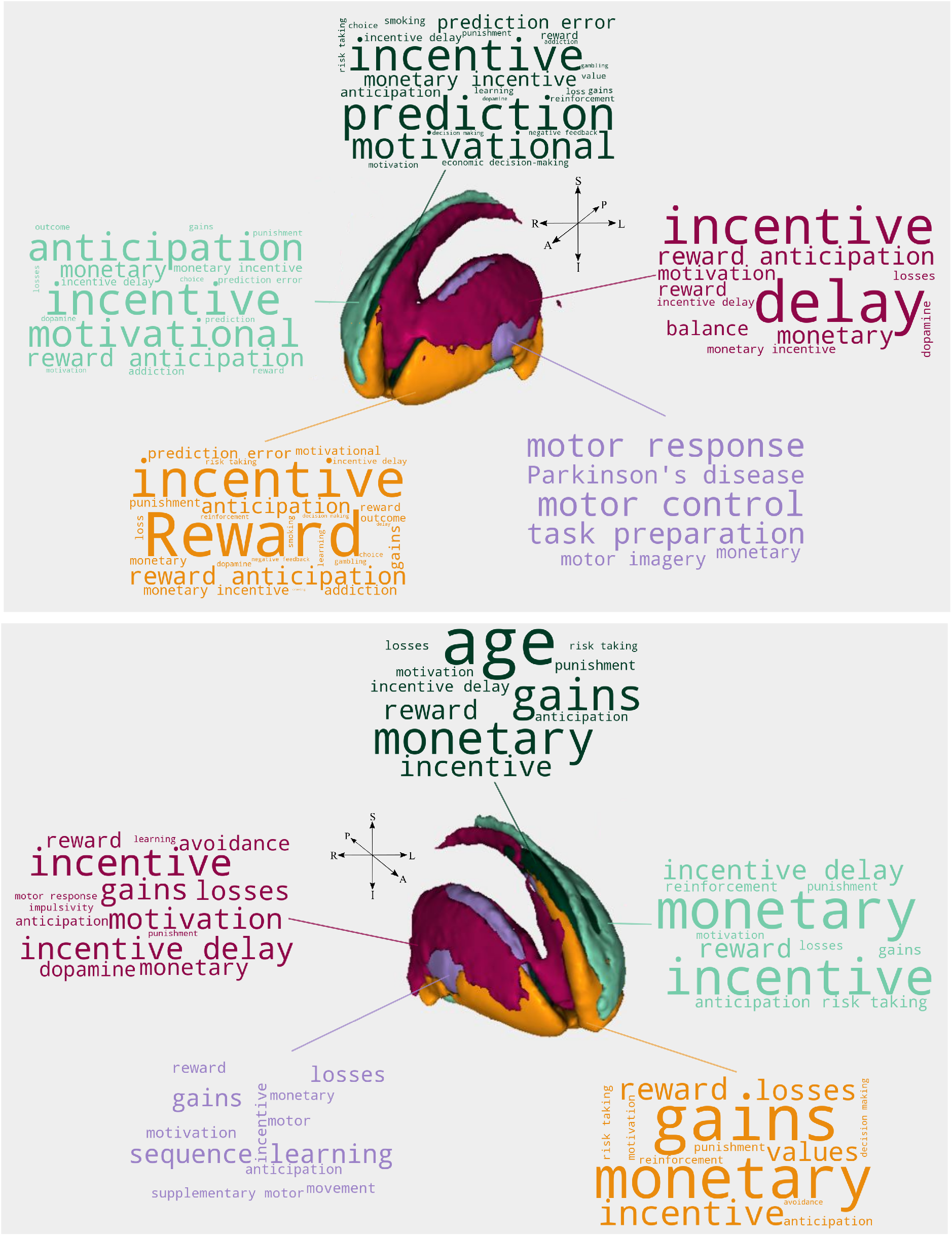
(Top) Left striatal components Neurosynth results.(Bottom) Right striatal components Neurosynth results. Here, the color of the words describe the components to which the posterior probability maps was related to (see ***Figure 3***C). The font of the word represents the Pearson correlation strength between the map of the component and the keyword related map from Neurosynth. Notice that the keywords’ font were not normalized across components. Hence, the keyword with the biggest font represents the term with the biggest correlation in that component and not in all components.

## Discussion

### Overview

We identified 5 spatially distinct microstructural components for the left and right striatum using OPNMF. We also found an increase in cluster stability when performing a multimodal decomposition rather than decomposing T1w/T2w, FA and MD data independently. By using brain-behaviour PLSC, we found four significant latent variables (two for each the left and right hemispheres) relating individual subject’s microstructural weightings in each component to behaviours and demographics. Finally, we also investigated how striatum clusters related to brain function using the Neurosynth database and ascertained some putative functional relationship of the specific clusters that we describe.

### Spatial striatal components and microstructure

Compared to other recent parcellations of the striatum, we notice that our multimodal clusters segregate across both the caudate and the putamen which have been observed in a recent (***Liu et al., 2020***) multi-modal parcellation of the striatum but not in other important data-driven parcellations (***Pauli et al., 2016***; ***Janssen et al., 2015***; ***Jung et al., 2014***). We also observe that the nucleus accumbens is encapsulated in its own cluster (component 4; orange) which is consistent with other striatum decompositions mentioned above.

Component 1 showed increased T1w/T2w in voxels corresponding to the dorsal putamen as well as some part of the posteromedial caudate. It has been observed that both FA and T1w/T2w are positively correlated with myelin density (***Uddin et al., 2019***). However, FA was shown to be a much stronger correlate of myelin content compared to T1w/T2w especially in subcortical grey matter structures (***Uddin et al., 2019***). Hence, the higher proportion of T1w/T2w compared to FA in component 1 might be attributed to another tissue microstructure property, like iron concentration(***Tardif et al., 2016***; ***Uddin et al., 2019***; ***Péran et al., 2009***).

Component 2 describes high FA compared to other metrics in voxels overlapping with a thin cluster along the anterior-posterior axis of lateral caudate and putamen. High FA might suggest a preferred fibre orientation in this region and myelination, although FA is sensitive to a wide range of cellular mechanism (***Tardif et al., 2016***; ***Uddin et al., 2019***). High T1w/T2w signal also suggests increased myelination in this region (***Uddin et al., 2019***), which combined with increased FA, could indicate the presence of fibre bundles. This may capture the anterior-posterior fibre organization in the caudate nucleus and inferior-superior myelinated fibre bundles between the caudate nucleus and globus pallidus through the internal capsule, which has recently been investigated using *in vivo* dMRI analyses and polarized light imaging in ***Kotz et al.*** (***2013***).

Component 4 included voxels overlapping with the nucleus accumbens structure as defined in ***Haber et al.*** (***1990***) and a thin cluster around the dorsal putamen. Component 4 describes increased FA and MD compared to T1w/T2w.

Component 5 is characterized by lower MD in some part of the inner anterior ventral caudate, the medial caudate body along the voxels of component 3 and some part of the ventral putamen. Decreased MD in these regions may suggest a denser tissue microstructure (***Beaulieu, 2002***; ***Sagi et al., 2012***).

The striatum has often been divided into functionally distinct regions based on corticostriatal inputs as there are no clear cytoarchitectonic parcellations of this structure. Tracing studies in non-human primates have identified a tripartite organization of the striatum based on structural connectivity to the cortex into the limbic region (ventral striatum), the association region (central striatum) and sensorimotor region (dorsolateral striatum) (***Haber et al., 1994***, ***1995***). Similar findings from tractography studies using diffusion MRI in humans have been observed in ***Draganski et al.*** (***2008***). The limbic region identified in (***Haber et al., 1994***, ***1995***, ***2006b***) overlaps with our component 4 (orange in ***Figure 3***) that also segregates the nucleus accumbens from the rest of the striatum. The association and sensorimotor regions from ***Haber et al. (1994***, 1995, 2006b) do not overlap as clearly with other components as our parcellation of the limbic region and component 4. However, we still see similarities between the association striatal regions and our fifth striatal component (dark green in ***Figure 3***), where both overlap with some part of the anterior caudate and anterior putamen. The somatosensory striatal region in ***Haber et al. (1995***, 2006b) corresponds the most to our component 1 (light purple in ***Figure 3***), comprised of the posterior putamen and posteromedial caudate.

Although our map does not exactly recapitulate this tripartite organization, we do see some similarities. This might suggest that some extrinsic structural connectivity properties of the striatum might be captured by the combination of intrinsic measures we used for our parcellation.

It is also known that the striatum contains two histochemically disinct compartments; the striosomes and matrix compartment (***Graybiel and Ragsdale, 1978***; ***Flaherty and Graybiel, 1994***; ***Holt et al., 1997***), that also differ in their input-output organization (***Gimenez-Amaya and Graybiel, 1991***; ***Eblen and Graybiel, 1995***). As the striosomes patches make up only 15% (***Brimblecombe and Cragg, 2017***) of the adult striatum and that these patches seems to be broadly distributed in the caudate and putamen (***Mikula et al., 2009***), it is not clear how this binary compartmentalization would affect our decomposition. Furthermore, current MRI protocols do not allow for the direct distinction between the striosome and matrix compartment (***Blood et al., 2018***) and it has yet been shown if and how the striosomes and matrix compartment affect the microstructural metrics derived from MRI that we used here.

### Individual-level variation in microstructure & behaviour

Microstructural components were also investigated at the individual level, where we assessed the relation between single-subject microstructure and behaviour. Using PLSC analysis, we identified two significant LVs for each the left and right striatum. Left LV1 and right LV1 displayed a similar pattern of increased MD across the left and right striatum correlated with young age, male sex and some measure of motor performance (increased strength and endurance and decreased dexterity). The inverse correlation between MD and age is consistent with evidence of decreased MD in early adulthood in deep grey matter structures (***Lebel et al., 2008***). The positive relationship between age and FA which is observed in left LV1 has also been established in ***Lebel et al.*** (***2008***), although here, this pattern is only observed in the left striatum. As for the behaviours, we note increased strength and decreased dexterity as well as an above-average performance of the Flanker task in both left and right LV1. The motor behaviour correlation pattern in the left and right LV1 is consistent with the sex effect observed in those LVs. Indeed, increased strength and endurance as well as decreased dexterity in males have been observed in those tasks before (***Hanten et al., 1999***; ***Bohannon et al., 2015***; ***Peters and Campagnaro, 1996***)

Left LV2 described a covariance mostly related to age and sex where young females exhibited a decrease in T1w/T2w across the left striatum and a decrease in FA in the putamen and the caudate nucleus (excluding the NA and the ‘outer rim’ of the putamen). The positive relationship between FA and age has been observed in previous studies (***Lebel et al., 2008***). Recent work has also identified a positive correlation between T1w/T2w and age during early adulthood, where a bilateral increase of T1w/T2w was observed in the striatum until a peak and subsequent decline at around 50 years old (***Tullo et al., 2019***). The left LV2 also displayed decreased strength and endurance. As for in the left and right LV1, we note that the motor pattern in left LV2 is also consistent with the sex effect in that LV.

The right LV2 displayed an inverse age-related pattern to the left LV2, where age correlated with increased FA in the entire striatum, increased T1w/T2w in the putamen and medial caudate along the ventricle, and decreased MD in the putamen. The positive relationship between age and FA in the right LV2 is consistent with findings in ***Lebel et al.*** (***2008***). The pattern of older age and increased T1w/T2w has also been observed in ***Tullo et al. (2019)***. In terms of behaviour and other demographics, this LV correlated with male sex, below-average performance in the Flanker task as well as increased strength and endurance.

As the female sample in this study has a slightly higher mean age than the males (mean female age = 29.01±3.62, mean male age = 27.71±3.67), the correlation patterns between the significant LVs with age and sex might be affected. For instance, a true correlation between an LV with age might also drive a correlation between the LV and sex or vice versa due to the previously noted bias in the sample. To investigate further the effect of sex in our LVs, we performed the same OPNMF followed by PLS on males and females independently. We found that for the left hemishpere, there was no significant difference between males and females in the striatum parcellation. Hence, we ran the PLS analysis for the left hemisphere without the sex as a variable and obtained similar LVs were the left LV1 mostly shows an effect of age and left LV2 shows an effect of the motor behaviours as seen in ***Figure 4***. As the right striatum parcellation was slightly different between males and females, we conducted the PLS analysis independently between males and females. We found that the microstructural partterns uncovered by the right LVs were different between males and females, probably due to the difference in the parcellation. However, the behavioural patterns were highly similar between the two groups. Indeed, the right LV1 for males and females shows mostly an effect of age and motor related behaviours while the right LV2 shows a stronger effect of impulsivity related behaviours, similar to what we show in ***Figure 4*** where the males and females were combined. More details on the sex specific analysis can be found in the supplement.

Moreover, relationships between brain structure and psychological traits using mass univariate approaches have been shown to have low replicability while there exist robust associations between brain structure and non-psychological traits such as age (***Masouleh et al., 2019***). This previously observed robust relationship between brain structure and age paired with the low variability in the HCP behavioural data might explain why we see such a strong effect of age and sex on our LVs compared to the other striatal related behaviours. In sum, we also note that the directionality of our PLSC results were as expected and we observed no laterality effects.

As discussed in another study from our group (***Patel et al., 2020***), the combination of OPNMF and PLS reduces the potential for false-positive as we are analyzing spatial components of voxels rather than performing univariate testing on every voxel. PLS is a multivariate technique that relates multiple variables simultaneously as opposed to multivariate testing, thus accounting for some difficulties encountered in univariate testing.

### Correlation with fMRI maps

The Neurosynth reverse-inference framework found multiple correlations between output components and posterior predictive maps associated with reward, incentive and decision-making related map, which are functions that have been attributed to the striatum in previous studies (***Stott and Redish, 2014***; ***Haber et al., 2006a***; ***van den Bos et al., 2014***; ***Pauli et al., 2016***; ***Jung et al., 2014***; ***Haber et al., 2006a***). Correlations with motor-related maps were stronger with putamen-related clusters (component 1, light purple), which is consistent with previous findings associating the putamen to somatosensory processes (***Arsalidou et al., 2013***; ***Pauli et al., 2016***). We found the strongest correlation with reward-related words in the bilateral component 4, which mostly overlaps with the nucleus accumbens. This is consistent with a recent finding (***Pauli et al., 2016***). However, the small size of our other components (component 2, 3 and 5) resulted in a major overlap between the components and the Neurosynth maps. Although all of the words are related to previously reported striatal fucntions, the component-map correlations are not particularly specific in components 2, 3 and 5.

We also note that the correlations uncovered by the Neurosynth framework are influenced by confirmation bias. For instance, studies that looked at reward or addiction related behaviours are more likely to mention the striatum or vice-versa as it has long been thought that such associations exists.

### Choice of parcellation

Striatal clustering Previous parcellations of the striatum have used a combination of heuristic and contrast-based definitions. In recent years, the increased quantity and quality of available MRI data have allowed for data-driven parcellations that rely on no *a priori* assumptions on striatal organization, overcoming the limitations of past parcellation schemes. To identify spatial striatal components, previous studies have used clustering techniques such as K-means clustering (***Pauli et al., 2016***; ***Parkes et al., 2017***; ***Jung et al., 2014***), and decomposition techniques such as PCA, ICA and probabilistic modelling, such as Gaussian mixture model (***Janssen et al., 2015***). Amongst the variety of possible parcellation schemes, one has to be careful in the selection of a clustering/decomposition algorithm as it depends heavily on the type of data and the aims of the study.

Although OPNMF has been shown to be mathematically equivalent to the K-means algorithm (***Eickhoff et al., 2018***), OPNMF was a suitable method for this study as we aimed to investigate inter-individual variability in the subjects’ weightings. ***Sotiras et al.*** (***2015***) showed that compared to other decomposition techniques (PCA and ICA), components captured by NMF seemed to reflect relevant biological processes related to age and were less prone to overfitting. The advantages of OPNMF interpretability have already been noted in previous studies (***Sotiras et al., 2015***; ***Patel et al., 2020***; ***Varikuti et al., 2018***). Here, we took advantage of the flexibility of NMF decomposition while capitalizing on a part-based representation of the striatum by adding the orthogonality constraint to NMF. We also note the data-driven symmetry in the component obtained in each hemisphere.

### Multimodal vs. unimodal

Although T1w/T2w, FA and MD are typically used in isolation, we hypothesized that since each of these measures has differential sensitivity to the underlying cellular anatomy but still some overlap in their range of sensitivities (i.e they are all sensitive to myelin) (***Glasser and Van Essen, 2011***; ***Tardif et al., 2016***; ***Tullo et al., 2019***), combining them would yield more robust parcels. Here, we note that the stability of the multimodal OPNMF decomposition was notably higher than the stability of the unimodal decomposition, which provides evidence for the benefit of integrating multiple metrics to construct a parcellation. There are multiple ways to obtain multimodal maps, however this is not a method typically employed in the literature. One way to obtain multi-modal maps is to superimpose all the parcellation schemes derived from one modality (***Eickhoff et al., 2018***). In this method, the final multimodal parcellation is based on the overlap of the voxels that had a similar cluster assignment in all the unimodal parcellation schemes (***Eickhoff et al., 2018***; ***Xia et al., 2017***; ***Wang et al., 2015***). Although such parcellation schemes provide useful confirmatory information, the voxels with ambiguous overlap between the distinct unimodal parcellation schemes were not necessarily included in the final map, which can lead to fragmented final multimodal parcellations (***Eickhoff et al., 2018***; ***Wang et al., 2015***).

OPNMF and other similar methods, such as PCA and ICA, try to overcome this limitation by integrating multiple modalities into the parcellation, making use of the confirmatory and complementary information provided by the multiple metrics.

### Limitations

An inherent limitation in this study is the lack of specificity regarding the underlying mechanism of structural and diffusion MRI derived metrics that we used. It is still not clear how specific aspects of tissue microstructure influence T1w/T2w, FA and MD. Other than myelin, the T1w and T2w signals are sensitive to the presence of macromolecules and iron concentration(***Tardif et al., 2016***; ***Uddin et al., 2019***). FA and MD are also sensitive to a wide range of additional cellular properties including axonal density and orientation, water in the tissue and the presence of different cell types (***Tardif et al., 2016***; ***Jones et al., 2013***). Although the combination of those microstructural metrics provides complementary and confirmatory information, it is still unclear what the underlying microstructure looks like in our identified striatal clusters. As with most non-invasive imaging studies, the resolution used in this study is subject to partial volume effects. Partial volume effects may affect metrics proportion in our striatal components, especially in components 2, 4 that are adjacent to major white matter tracts which might be contributing to the in-crease of FA. Partial volume effects may also play a role in the high proportion of MD in component 3 as it is adjacent to the the anterior horn of the lateral ventricle.

## Conclusion

In this work, we used a combination of three microstructural metrics to construct a part-based decomposition of the human striatum in a healthy population using non-negative matrix factorization. By using the stability and accuracy of OPNMF decomposition, we identified 5 spatially distinct microstructural patterns for the left and right striatum separately. Then, we used partial least squares correlation to link inter-individual variation in the striatal components to selected behaviours and demographics. Our findings suggest distinct microstructural patterns in the human striatum that relate mostly to demographics. Our work also highlights the gain in clusters’ stability when using multimodal versus unimodal metrics. We note that the identified striatal components are associated with complex patterns of microstructure and behavioural variation. Further, the striatal components appear to be functionally relevant.

This work can serve as a template for examining how one can investigate subject-level variation that links brain and behaviour across numerous brain imaging measures. This may, in turn, allow for more specific interpretations of brain imaging findings that improve our mechanistic insights on brain-behaviour relationships. Further, this work could be applied in future studies of brain development and in the context of neuropsychiatric disorders to parse heterogeneity.

## Acknowledgments

Data were provided [in part] by the Human Connectome Project, WU-Minn Consortium (Principal Investigators: David Van Essen and Kamil Ugurbil; 1U54MH091657) funded by the 16 NIH Institutes and Centers that support the NIH Blueprint for Neuroscience Research; and by the McDonnell Center for Systems Neuroscience at Washington University.

## References

Albin RL, Young AB, Penney JB. The functional anatomy of basal ganglia disorders. . 1989;.

Alexander AL, Lee JE, Lazar M, Field AS. Diffusion Tensor Imaging of the Brain. Neurotherapeutics. 2007; 4(3):316 – 329. http://www.sciencedirect.com/science/article/pii/S1933721307000955, doi: https://doi.org/10.1016/j.nurt.2007.05.011, advances in Neuroimaging/Neuroethics.

Arsalidou M, Duerden EG, Taylor MJ. The centre of the brain: Topographical model of motor, cognitive, affective, and somatosensory functions of the basal ganglia. Human Brain Mapping. 2013; 34(11):3031–3054. https://onlinelibrary.wiley.com/doi/abs/10.1002/hbm.22124 doi: https://doi.org/10.1002/hbm.22124.

Avants BB, Yushkevich P, Pluta J, Minkoff D, Korczykowski M, Detre J, Gee JC. The optimal template effect in hippocampus studies of diseased populations. Neuroimage. 2010; 49(3):2457–2466.

Basser PJ, Mattiello J, LeBihan D. Estimation of the effective self-diffusion tensor from the NMR spin echo. J Magn Reson B. 1994; 103(3):247–54. doi: 10.1006/jmrb.1994.1037.

Basser PJ, Mattiello J, LeBihan D. MR diffusion tensor spectroscopy and imaging. Biophys J. 1994; 66(1):259–67. doi: 10.1016/s0006-3495(94)80775-1.

Beaulieu C. The basis of anisotropic water diffusion in the nervous system–a technical review. NMR in Biomedicine: An International Journal Devoted to the Development and Application of Magnetic Resonance In Vivo. 2002; 15(7-8):435–455.

Blood AJ, Waugh JL, Münte TF, Heldmann M, Domingo A, Klein C, Breiter HC, Lee LV, Rosales RL, Brüggemann N. Increased insula-putamen connectivity in X-linked dystonia-parkinsonism. NeuroImage: Clinical. 2018; 17:835–846. https://www.sciencedirect.com/science/article/pii/S2213158217302656, doi: https://doi.org/10.1016/j.nicl.2017.10.025.

Bohannon RW, Wang YC, Gershon RC. Two-minute walk test performance by adults 18 to 85 years: normative values, reliability, and responsiveness. Archives of physical medicine and rehabilitation. 2015; 96(3):472–477.

van den Bos W, Rodriguez CA, Schweitzer JB, McClure SM. Connectivity Strength of Dissociable Striatal Tracts Predict Individual Differences in Temporal Discounting. The Journal of Neuroscience. 2014; 34(31):10298. http://www.jneurosci.org/content/34/31/10298.abstracthttps://www.ncbi.nlm.nih.gov/pmc/articles/PMC4577570/pdf/zns10298.pdf , doi: 10.1523/JNEUROSCI.4105-13.2014.

Boutsidis C, Gallopoulos E. SVD based initialization: A head start for nonnegative matrix factorization. Pattern Recognition. 2008; 41(4):1350–1362. http://www.sciencedirect.com/science/article/pii/S0031320307004359, doi: https://doi.org/10.1016/j.patcog.2007.09.010.

Brimblecombe KR, Cragg SJ. The striosome and matrix compartments of the striatum: a path through the labyrinth from neurochemistry toward function. ACS chemical neuroscience. 2017; 8(2):235–242. https://pubs.acs.org/doi/pdf/10.1021/acschemneuro.6b00333.

Buckholtz JW, Treadway MT, Cowan RL, Woodward ND, Li R, Ansari MS, Baldwin RM, Schwartzman AN, Shelby ES, Smith CE, Kessler RM, Zald DH. Dopaminergic Network Differences in Human Impulsivity. Science. 2010; 329(5991):532–532. https://science.sciencemag.org/content/329/5991/532, doi: 10.1126/science.1185778.

Burrer A, Caravaggio F, Manoliu A, Plitman E, Gütter K, Habermeyer B, Stämpfli P, Abivardi A, Schmidt A, Borgwardt S, Chakravarty M, Lepage M, Dagher A, Graff-Guerrero A, Seifritz E, Kaiser S, Kirschner M. Apathy is not associated with reduced ventral striatal volume in patients with schizophrenia. Schizophr Res. 2020; 223:279–288. doi: 10.1016/j.schres.2020.08.018.

Caravaggio F, Plavén-Sigray P, Matheson GJ, Plitman E, Chakravarty MM, Borg J, Graff-Guerrero A, Cervenka S. Trait impulsivity is not related to post-commissural putamen volumes: A replication study in healthy men. PLoS One. 2018; 13(12):e0209584. doi: 10.1371/journal.pone.0209584.

Chakravarty MM, Bertrand G, Hodge CP, Sadikot AF, Collins DL. The creation of a brain atlas for image guided neurosurgery using serial histological data. Neuroimage. 2006; 30(2):359–76. doi: 10.1016/j.neuroimage.2005.09.041.

Chakravarty MM, Rapoport JL, Giedd JN, Raznahan A, Shaw P, Collins DL, Lerch JP, Gog-tay N. Striatal shape abnormalities as novel neurodevelopmental endophenotypes in schizophrenia: a longitudinal study. Human brain mapping. 2015; 36(4):1458–1469. http://europepmc.org/abstract/MED/25504933https://doi.org/10.1002/hbm.22715https://europepmc.org/articles/PMC6869651https://europepmc.org/articles/PMC6869651?pdf=render, doi: 10.1002/hbm.22715.

Chakravarty MM, Steadman P, van Eede MC, Calcott RD, Gu V, Shaw P, Raznahan A, Collins DL, Lerch JP. Performing label-fusion-based segmentation using multiple automatically generated templates. Human brain mapping. 2013; 34(10):2635–2654.

Chang LJ, Yarkoni T, Khaw MW, Sanfey AG. Decoding the Role of the Insula in Human Cognition: Functional Parcellation and Large-Scale Reverse Inference. Cerebral Cortex. 2012 03; 23(3):739–749. https://doi.org/10.1093/cercor/bhs065, doi: a10.1093/cercor/bhs065.

Choi EY, Yeo BT, Buckner RL. The organization of the human striatum estimated by intrinsic functional connectivity. Journal of neurophysiology. 2012; 108(8):2242–2263.

Dalley JW, Mar AC, Economidou D, Robbins TW. Neurobehavioral mechanisms of impulsivity: Fronto-striatal systems and functional neurochemistry. Pharmacology Biochemistry and Behavior. 2008; 90(2):250 – 260. http://www.sciencedirect.com/science/article/pii/S0091305707003826, doi: https://doi.org/10.1016/j.pbb.2007.12.021, microdialysis: recent developments.

Delong MR, Crutcher MD, Georgopoulos AP. Relations between movement and single cell discharge in the substantia nigra of the behaving monkey. Journal of Neuroscience. 1983; 3(8):1599–1606.

Draganski B, Kherif F, Klöppel S, Cook PA, Alexander DC, Parker GJM, Deichmann R, Ashburner J, Frackowiak RSJ. Evidence for Segregated and Integrative Connectivity Patterns in the Human Basal Ganglia. The Journal of Neuroscience. 2008; 28(28):7143. http://www.jneurosci.org/content/28/28/7143.abstracthttps://www.ncbi.nlm.nih.gov/pmc/articles/PMC6670486/pdf/zns7143.pdf, doi: 10.1523/JNEUROSCI.1486-08.2008.

Eblen F, Graybiel A. Highly restricted origin of prefrontal cortical inputs to striosomes in the macaque monkey. Journal of Neuroscience. 1995; 15(9):5999–6013. https://www.jneurosci.org/content/15/9/5999, doi: 10.1523/JNEUROSCI.15-09-05999.1995.

Eickhoff SB, Yeo BTT, Genon S. Imaging-based parcellations of the human brain. Nature Reviews Neuroscience. 2018; 19(11):672–686. https://doi.org/10.1038/s41583-018-0071-7https://www.nature.com/articles/s41583-018-0071-7.pdf, doi: 10.1038/s41583-018-0071-7.

Estle SJ, Green L, Myerson J, Holt DD. Differential effects of amount on temporal and probability discounting of gains and losses. Memory & Cognition. 2006; 34(4):914–928.

Flaherty A, Graybiel AM. Input-output organization of the sensorimotor striatum in the squirrel monkey. Journal of Neuroscience. 1994; 14(2):599–610.

Gimenez-Amaya J, Graybiel A. Modular organization of projection neurons in the matrix compartment of the primate striatum. Journal of Neuroscience. 1991; 11(3):779–791. https://www.jneurosci.org/content/11/3/779, doi: 10.1523/JNEUROSCI.11-03-00779.1991.

Glasser MF, Coalson TS, Robinson EC, Hacker CD, Harwell J, Yacoub E, Ugurbil K, Andersson J, Beckmann CF, Jenkinson M, Smith SM, Van Essen DC. A multi-modal parcellation of human cerebral cortex. Nature. 2016; 536(7615):171–178. https://doi.org/10.1038/nature18933, doi: 10.1038/nature18933.

Glasser MF, Sotiropoulos SN, Wilson JA, Coalson TS, Fischl B, Andersson JL, Xu J, Jbabdi S, Webster M, Polimeni JR, Van Essen DC, Jenkinson M. The minimal preprocessing pipelines for the Human Connectome Project. NeuroImage. 2013; 80:105 – 124. http://www.sciencedirect.com/science/article/pii/S1053811913005053, doi: https://doi.org/10.1016/j.neuroimage.2013.04.127, mapping the Connectome.

Glasser MF, Van Essen DC. Mapping Human Cortical Areas In Vivo Based on Myelin Content as Revealed by T1- and T2-Weighted MRI. Journal of Neuroscience. 2011; 31(32):11597–11616. https://www.jneurosci.org/content/31/32/11597, doi: 10.1523/JNEUROSCI.2180-11.2011.

Graybiel AM, Grafton ST. The striatum: where skills and habits meet. Cold Spring Harbor perspectives in biology. 2015; 7(8):a021691. https://www.ncbi.nlm.nih.gov/pmc/articles/PMC4526748/pdf/cshperspect-LNM-a021691.pdf.

Graybiel AM, Ragsdale CW. Histochemically distinct compartments in the striatum of human, monkeys, and cat demonstrated by acetylthiocholinesterase staining. Proceedings of the National Academy of Sciences. 1978; 75(11):5723–5726.

Graybiel AM, Rauch SL. Toward a neurobiology of obsessive-compulsive disorder. Neuron. 2000; 28(2):343–347.

Green L, Myerson J, Shah AK, Estle SJ, Holt DD. Do adjusting-amount and adjusting-delay procedures produce equivalent estimates of subjective value in pigeons? Journal of the experimental analysis of behavior. 2007; 87(3):337–347.

Haber SN, Lynd E, Klein C, Groenewegen HJ. Topographic organization of the ventral striatal efferent projections in the rhesus monkey: an anterograde tracing study. J Comp Neurol. 1990; 293(2):282–98. doi: 10.1002/cne.902930210.

Haber SN, Kim KS, Mailly P, Calzavara R. Reward-related cortical inputs define a large striatal region in primates that interface with associative cortical connections, providing a substrate for incentive-based learning. Journal of Neuroscience. 2006; 26(32):8368–8376. https://www.ncbi.nlm.nih.gov/pmc/articles/PMC6673798/pdf/zns8368.pdf.

Haber SN, Kim KS, Mailly P, Calzavara R. Reward-related cortical inputs define a large striatal region in primates that interface with associative cortical connections, providing a substrate for incentive-based learning. The Journal of neuroscience : the official journal of the Society for Neuroscience. 2006; 26(32):8368–8376. https://pubmed.ncbi.nlm.nih.gov/16899732https://www.ncbi.nlm.nih.gov/pmc/articles/PMC6673798/, doi: 10.1523/JNEUROSCI.0271-06.2006.

Haber SN, Kunishio K, Mizobuchi M, Lynd-Balta E. The orbital and medial prefrontal circuit through the primate basal ganglia. Journal of Neuroscience. 1995; 15(7):4851–4867.

Haber SN, Lynd-Balta E, Spooren WP. Integrative aspects of basal ganglia circuitry. In: The basal ganglia IV Springer; 1994.p. 71–80.

Hacker CD, Perlmutter JS, Criswell SR, Ances BM, Snyder AZ. Resting state functional connectivity of the striatum in Parkinson’s disease. Brain. 2012 11; 135(12):3699–3711. https://doi.org/10.1093/brain/aws281, doi: 10.1093/brain/aws281.

Halko N, Martinsson PG, Tropp JA. Finding structure with randomness: Probabilistic algorithms for constructing approximate matrix decompositions. SIAM review. 2011; 53(2):217–288.

Hanten WP, Chen WY, Austin AA, Brooks RE, Carter HC, Law CA, Morgan MK, Sanders DJ, Swan CA, Vanderslice AL. Maximum grip strength in normal subjects from 20 to 64 years of age. Journal of hand therapy. 1999; 12(3):193–200.

Hare TA, Tottenham N, Davidson MC, Glover GH, Casey BJ. Contributions of amygdala and striatal activity in emotion regulation. Biological Psychiatry. 2005; 57(6):624–632. http://www.sciencedirect.com/science/article/pii/S0006322304013812https://www.biologicalpsychiatryjournal.com/article/S0006-3223(04)01381-2/fulltext, doi: https://doi.org/10.1016/j.biopsych.2004.12.038.

Hariri AR, Brown SM, Williamson DE, Flory JD, de Wit H, Manuck SB. Preference for Immediate over Delayed Rewards Is Associated with Magnitude of Ventral Striatal Activity. Journal of Neuroscience. 2006; 26(51):13213–13217. https://www.jneurosci.org/content/26/51/13213, doi: 10.1523/JNEUROSCI.3446-06.2006.

Holt DJ, Graybiel AM, Saper CB. Neurochemical architecture of the human striatum. Journal of Comparative Neurology. 1997; 384(1):1–25.

Janssen RJ, Jylänki P, Kessels RPC, van Gerven MAJ. Probabilistic model-based functional parcellation reveals a robust, fine-grained subdivision of the striatum. NeuroImage. 2015; 119:398 – 405. http://www.sciencedirect.com/science/article/pii/S1053811915005893, doi: https://doi.org/10.1016/j.neuroimage.2015.06.084.

Jones DK, Knösche TR, Turner R. White matter integrity, fiber count, and other fallacies: The do’s and don’ts of diffusion MRI. NeuroImage. 2013; 73:239–254. http://www.sciencedirect.com/science/article/pii/S1053811912007306https://www.sciencedirect.com/science/article/abs/pii/S1053811912007306?via%3Dihub, doi: https://doi.org/10.1016/j.neuroimage.2012.06.081.

Jung WH, Jang JH, Park JW, Kim E, Goo EH, Im OS, Kwon JS. Unravelling the intrinsic functional organization of the human striatum: a parcellation and connectivity study based on resting-state FMRI. PloS one. 2014; 9(9):e106768. https://www.ncbi.nlm.nih.gov/pmc/articles/PMC4159235/pdf/pone.0106768.pdf.

Kotz SA, Anwander A, Axer H, Knösche TR. Beyond Cytoarchitectonics: The Internal and External Connectivity Structure of the Caudate Nucleus. PLOS ONE. 2013; 8(7):e70141. https://doi.org/10.1371/journal.pone.0070141, doi: 10.1371/journal.pone.0070141.

Krishnan A, Williams LJ, McIntosh AR, Abdi H. Partial Least Squares (PLS) methods for neuroimaging: A tutorial and review. NeuroImage. 2011; 56(2):455 – 475. http://www.sciencedirect.com/science/article/pii/S1053811910010074, doi: https://doi.org/10.1016/j.neuroimage.2010.07.034, multivariate Decoding and Brain Reading.

Lebel C, Walker L, Leemans A, Phillips L, Beaulieu C. Microstructural maturation of the human brain from childhood to adulthood. Neuroimage. 2008; 40(3):1044–1055.

Leh SE, Ptito A, Chakravarty MM, Strafella AP. Fronto-striatal connections in the human brain: a probabilistic diffusion tractography study. Neuroscience letters. 2007; 419(2):113–118. https://pubmed.ncbi.nlm.nih.gov/17485168https://www.ncbi.nlm.nih.gov/pmc/articles/PMC5114128/, doi: 10.1016/j.neulet.2007.04.049.

Lehéricy S, Ducros M, Van De Moortele PF, Francois C, Thivard L, Poupon C, Swindale N, Ugurbil K, Kim DS. Diffusion tensor fiber tracking shows distinct corticostriatal circuits in humans. Annals of Neurology. 2004; 55(4):522–529. https://onlinelibrary.wiley.com/doi/abs/10.1002/ana.20030, doi: https://doi.org/10.1002/ana.20030.

Li Y, Yuan K, Cai C, Feng D, Yin J, Bi Y, Shi S, Yu D, Jin C, von Deneen KM, Qin W, Tian J. Reduced frontal cortical thickness and increased caudate volume within fronto-striatal circuits in young adult smokers. Drug and Alcohol Dependence. 2015; 151:211–219. http://www.sciencedirect.com/science/article/pii/S0376871615001799https://www.sciencedirect.com/science/article/abs/pii/S0376871615001799?via%3Dihub, doi: https://doi.org/10.1016/j.drugalcdep.2015.03.023.

Liu X, Eickhoff SB, Hoffstaedter F, Genon S, Caspers S, Reetz K, Dogan I, Eickhoff CR, Chen J, Caspers J, Reuter N, Mathys C, Aleman A, Jardri R, Riedl V, Sommer IE, Patil KR. Joint Multi-modal Parcellation of the Human Striatum: Functions and Clinical Relevance. Neuroscience Bulletin. 2020; 36(10):1123–1136. https://doi.org/10.1007/s12264-020-00543-1, doi: 10.1007/s12264-020-00543-1.

Marquand AF, Haak KV, Beckmann CF. Functional corticostriatal connection topographies predict goal directed behaviour in humans. Nature human behaviour. 2017; 1(8):0146–0146. https://pubmed.ncbi.nlm.nih.gov/28804783https://www.ncbi.nlm.nih.gov/pmc/articles/PMC5549843/, doi: 10.1038/s41562-017-0146.

Masouleh SK, Eickhoff SB, Hoffstaedter F, Genon S, Initiative ADN, et al. Empirical examination of the replicability of associations between brain structure and psychological variables. Elife. 2019; 8:e43464.

McIntosh AR, Lobaugh NJ. Partial least squares analysis of neuroimaging data: applications and advances. NeuroImage. 2004; 23:S250 – S263. http://www.sciencedirect.com/science/article/pii/S1053811904003866, doi: https://doi.org/10.1016/j.neuroimage.2004.07.020, mathematics in Brain Imaging.

Mikula S, Parrish SK, Trimmer JS, Jones EG. Complete 3D visualization of primate striosomes by KChIP1 immunostaining. Journal of Comparative Neurology. 2009; 514(5):507–517. https://onlinelibrary.wiley.com/doi/abs/10.1002/cne.22051, doi: https://doi.org/10.1002/cne.22051.

Milad MR, Rauch SL. Obsessive-compulsive disorder: beyond segregated cortico-striatal pathways. Trends in Cognitive Sciences. 2012; 16(1):43 – 51. http://www.sciencedirect.com/science/article/pii/S1364661311002361, doi: https://doi.org/10.1016/j.tics.2011.11.003, special Issue: Cognition in Neuropsychiatric Disorders.

Mink JW. The basal ganglia: focused selection and inhibition of competing motor programs. Progress in neurobiology. 1996; 50(4):381–425.

Myerson J, Green L, Warusawitharana M. Area under the curve as a measure of discounting. Journal of the experimental analysis of behavior. 2001; 76(2):235–243.

Parkes L, Fulcher BD, Yücel M, Fornito A. Transcriptional signatures of connectomic subregions of the human striatum. Genes, Brain and Behavior. 2017; 16(7):647–663. https://onlinelibrary.wiley.com/doi/abs/10.1111/gbb.12386, doi: https://doi.org/10.1111/gbb.12386.

Patel R, Steele CJ, Chen AGX, Patel S, Devenyi GA, Germann J, Tardif CL, Chakravarty MM. Investigating microstructural variation in the human hippocampus using non-negative matrix factorization. NeuroImage. 2020; 207:116348. http://www.sciencedirect.com/science/article/pii/S1053811919309395, doi: https://doi.org/10.1016/j.neuroimage.2019.116348.

Pauli WM, O’Reilly RC, Yarkoni T, Wager TD. Regional specialization within the human striatum for diverse psychological functions. Proceedings of the National Academy of Sciences. 2016; 113(7):1907–1912. https://www.pnas.org/content/113/7/1907, doi: 10.1073/pnas.1507610113.

Peters M, Campagnaro P. Do women really excel over men in manual dexterity? Journal of Experimental Psychology: Human Perception and Performance. 1996; 22(5):1107.

Péran P, Cherubini A, Luccichenti G, Hagberg G, Démonet JF, Rascol O, Celsis P, Caltagirone C, Spalletta G, Sabatini U. Volume and iron content in basal ganglia and thalamus. Human brain mapping. 2009; 30(8):2667–2675. https://pubmed.ncbi.nlm.nih.gov/19172651https://www.ncbi.nlm.nih.gov/pmc/articles/PMC6871035/, doi: 10.1002/hbm.20698.

Rolls E. Neurophysiology and cognitive functions of the striatum. Revue neurologique. 1994; 150 8-9:648–60.

Rosenblatt A, Leroi I. Neuropsychiatry of Huntington’s Disease and Other Basal Ganglia Disorders. Psychosomatics. 2000; 41(1):24 – 30. http://www.sciencedirect.com/science/article/pii/S0033318200711704, doi: https://doi.org/10.1016/S0033-3182(00)71170-4.

Sagi Y, Tavor I, Hofstetter S, Tzur-Moryosef S, Blumenfeld-Katzir T, Assaf Y. Learning in the Fast Lane: New Insights into Neuroplasticity. Neuron. 2012; 73(6):1195–1203. http://www.sciencedirect.com/science/article/pii/S089662731200178X, doi: https://doi.org/10.1016/j.neuron.2012.01.025.

Schouppe N, Demanet J, Boehler CN, Ridderinkhof KR, Notebaert W. The Role of the Striatum in Effort-Based Decision-Making in the Absence of Reward. Journal of Neuroscience. 2014; 34(6):2148–2154. https://www.jneurosci.org/content/34/6/2148, doi: 10.1523/JNEUROSCI.1214-13.2014.

Schuetze M, Park MT, Cho IY, MacMaster FP, Chakravarty MM, Bray SL. Morphological Alterations in the Thalamus, Striatum, and Pallidum in Autism Spectrum Disorder. Neuropsychopharmacology. 2016; 41(11):2627–37. doi: 10.1038/npp.2016.64.

Shaw P, Sharp W, Sudre G, Wharton A, Greenstein D, Raznahan A, Evans A, Chakravarty MM, Lerch JP, Rapoport J. Subcortical and cortical morphological anomalies as an endophenotype in obsessive-compulsive disorder. Mol Psychiatry. 2015; 20(2):224–31. doi: 10.1038/mp.2014.3.

Sotiras A, Resnick SM, Davatzikos C. Finding imaging patterns of structural covariance via Non-Negative Matrix Factorization. NeuroImage. 2015; 108:1–16. https://pubmed.ncbi.nlm.nih.gov/25497684https://www.ncbi.nlm.nih.gov/pmc/articles/PMC4357179/https://www.ncbi.nlm.nih.gov/pmc/articles/PMC4357179/pdf/nihms649302.pdf, doi: 10.1016/j.neuroimage.2014.11.045.

Stott JJ, Redish AD. A functional difference in information processing between orbitofrontal cortex and ventral striatum during decision-making behaviour. Philosophical Transactions of the Royal Society B: Biological Sciences. 2014; 369(1655):20130472. https://royalsocietypublishing.org/doi/abs/10.1098/rstb.2013.0472, doi: doi:10.1098/rstb.2013.0472.

Tardif CL, Gauthier CJ, Steele CJ, Bazin PL, Schäfer A, Schaefer A, Turner R, Villringer A. Advanced MRI techniques to improve our understanding of experience-induced neuroplasticity. NeuroImage. 2016; 131:55–72. http://www.sciencedirect.com/science/article/pii/S1053811915007661https://www.sciencedirect.com/science/article/abs/pii/S1053811915007661?via%3Dihub, doi: https://doi.org/10.1016/j.neuroimage.2015.08.047.

Tournier JD, Calamante F, Connelly A. MRtrix: diffusion tractography in crossing fiber regions. International journal of imaging systems and technology. 2012; 22(1):53–66.

Tullo S, Devenyi GA, Patel R, Park MTM, Collins DL, Chakravarty MM. Warping an atlas derived from serial histology to 5 high-resolution MRIs. Scientific data. 2018; 5:180107.

Tullo S, Patel R, Devenyi GA, Salaciak A, Bedford SA, Farzin S, Wlodarski N, Tardif CL, Group PAR, Breitner JCS, Chakravarty MM. MR-based age-related effects on the striatum, globus pallidus, and thalamus in healthy individuals across the adult lifespan. Human brain mapping. 2019; 40(18):5269–5288. https://pubmed.ncbi.nlm.nih.gov/31452289https://www.ncbi.nlm.nih.gov/pmc/articles/PMC6864890/, doi: 10.1002/hbm.24771.

Tziortzi AC, Haber SN, Searle GE, Tsoumpas C, Long CJ, Shotbolt P, Douaud G, Jbabdi S, Behrens TEJ, Rabiner EA, Jenkinson M, Gunn RN. Connectivity-Based Functional Analysis of Dopamine Release in the Striatum Using Diffusion-Weighted MRI and Positron Emission Tomography. Cerebral Cortex. 2014; 24(5):1165–1177. https://doi.org/10.1093/cercor/bhs397https://www.ncbi.nlm.nih.gov/pmc/articles/PMC3977617/pdf/bhs397.pdf, doi: 10.1093/cercor/bhs397.

Uddin MN, Figley TD, Solar KG, Shatil AS, Figley CR. Comparisons between multi-component myelin water fraction, T1w/T2w ratio, and diffusion tensor imaging measures in healthy human brain structures. Scientific Reports. 2019; 9(1):2500. https://doi.org/10.1038/s41598-019-39199-x https://www.ncbi.nlm.nih.gov/pmc/articles/PMC6384876/pdf/41598_2019_Article_39199.pdf, doi: 10.1038/s41598-019-39199-x.

Van Essen DC, Smith SM, Barch DM, Behrens TEJ, Yacoub E, Ugurbil K. The WU-Minn Human Connectome Project: An overview. NeuroImage. 2013; 80:62–79. http://www.sciencedirect.com/science/article/pii/S1053811913005351, doi: https://doi.org/10.1016/j.neuroimage.2013.05.041.

Varikuti DP, Genon S, Sotiras A, Schwender H, Hoffstaedter F, Patil KR, Jockwitz C, Caspers S, Moebus S, Amunts K. Evaluation of non-negative matrix factorization of grey matter in age prediction. Neuroimage. 2018; 173:394–410. https://www.ncbi.nlm.nih.gov/pmc/articles/PMC5911196/pdf/nihms952495.pdf.

Veraart J, Sijbers J, Sunaert S, Leemans A, Jeurissen B. Weighted linear least squares estimation of diffusion MRI parameters: strengths, limitations, and pitfalls. Neuroimage. 2013; 81:335–346. doi: 10.1016/j.neuroimage.2013.05.028.

Vink M, Kahn RS, Raemaekers M, van den Heuvel M, Boersma M, Ramsey NF. Function of striatum beyond inhibition and execution of motor responses. Human Brain Mapping. 2005; 25(3):336–344. https://doi.org/10.1002/hbm.20111https://www.ncbi.nlm.nih.gov/pmc/articles/PMC6871687/pdf/HBM-25-336.pdf, doi: 10.1002/hbm.20111.

Wang J, Yang Y, Fan L, Xu J, Li C, Liu Y, Fox PT, Eickhoff SB, Yu C, Jiang T. Convergent functional architecture of the superior parietal lobule unraveled with multimodal neuroimaging approaches. Human brain mapping. 2015; 36(1):238–257.

Westin CF, Peled S, Gudbjartsson H, Kikinis R, Jolesz FA. Geometrical Diffusion Measures for MRI from Tensor Basis Analysis. In: ISMRM’97 Vancouver Canada; 1997. p. 1742.

Xia X, Fan L, Cheng C, Eickhoff SB, Chen J, Li H, Jiang T. Multimodal connectivity-based parcellation reveals a shell-core dichotomy of the human nucleus accumbens. Human Brain Mapping. 2017; 38(8):3878–3898. https://www.ncbi.nlm.nih.gov/pmc/articles/PMC5685173/pdf/HBM-38-3878.pdf, doi: 10.1002/hbm.23636.

Yager LM, Garcia AF, Wunsch AM, Ferguson SM. The ins and outs of the striatum: Role in drug addiction. Neuroscience. 2015; 301:529 – 541. http://www.sciencedirect.com/science/article/pii/S0306452215005746, doi: https://doi.org/10.1016/j.neuroscience.2015.06.033.

Yang Z, Oja E. Linear and Nonlinear Projective Nonnegative Matrix Factorization. IEEE Transactions on Neural Networks. 2010; 21(5):734–749. doi: 10.1109/TNN.2010.2041361.

Yarkoni T, Poldrack RA, Nichols TE, Van Essen DC, Wager TD. Large-scale automated synthesis of human functional neuroimaging data. Nature Methods. 2011; 8(8):665–670. https://doi.org/10.1038/nmeth.1635, doi: 10.1038/nmeth.1635.

Zeighami Y, Fereshtehnejad SM, Dadar M, Collins DL, Postuma RB, Mišić B, Dagher A. A clinicalanatomical signature of Parkinson’s disease identified with partial least squares and magnetic resonance imaging. NeuroImage. 2019; 190:69 – 78. http://www.sciencedirect.com/science/article/pii/S1053811917310741, doi: https://doi.org/10.1016/j.neuroimage.2017.12.050, mapping diseased brains.

