## Supplemental Material for "Microstructural variation in the human striatum using non-negative matrix factorization"

### Supplement

#### NMF Background

Let $X$ (m x n) be the input matrix, then the basis of matrix factorization consists of decomposing $X$ into two matrices, the matrix of component $W$ (m x K) and the weight coefficient matrix $H$ (k x n) which represents the strength of each component such that $X = WH$ (***Sotiras et al., 2015***). In NMF, both the components and the weights must be positive. The matrices W and H are the non-negative matrices that minimize the Frobenius norm $\|X - WH\|_F^2$ , thus $W$ and $H$ are the best estimates of $X$ (***Sotiras et al., 2015***). As this decomposition is non-negative, the input can be interpreted as an addition of positive components it is easier to interpret in term of brain imaging as all of the metrics that we use have positive values (***Sotiras et al., 2015***). Now adding the projection constraint (PNMF) on regular NMF, we assume that the loading matrix is a projection of the data matrix, i.e $H = W^T X$ (***Sotiras et al., 2015***). In OPNMF we further assume that the component matrix $W$ is orthonormal. Hence, OPNMF solves the minimization problem

$$\min_W \|X - WW^T X\|_F^2 \text{ where } WW^T = I \text{ and } W \geq 0 \text{ (Sotiras et al., 2015)} \quad (1)$$

Here, OPNMF to extract non-overlapping components from neuroimaging data, whereas PNMF or traditional NMF can result in a softer partitioning. To further ensure non-overlapping components, OPNMF components matrix $W$ can be initialized using non-negative double singular value decomposition (SVD) (***Boutsidis and Gallopoulos, 2008***; ***Patel et al., 2020***). Then W is iteratively updated as follows

$$W'_{ij} = W_{ij} \frac{(XX^T W)_{ij}}{(WW^T XX^T W)_{ij}} \quad (2)$$

Where $i$ is the voxel and $j$ the specific component (***Patel et al., 2020***; ***Varikuti et al., 2018***). The weight matrix $H$ can then be computed by

$$H = W^T X \quad (3)$$

The components in $W$ identified by OPNMF can be interpreted as parts of the original data, while the weight in $H$ represents the loading of each sample on a component to reconstruct the original data (***Patel et al., 2020***).

#### Partial Least Squares Background

To associate the selected behaviours to the subject's metrics proportion, we used Partial Least Squares (PLSC). PLSC is a multivariate statistical technique that analyses the association between two sets of high-dimensional variables, namely brain-derived and behavioural variables in the context of the current study (***Krishnan et al., 2011***; ***Zeighami et al., 2019***; ***McIntosh and Lobaugh, 2004***; ***Patel et al., 2020***). Let the matrix $X$ contains the neuroimaging or brain-derived variable and the matrix $Y$ containing the behavioural variables. Both $X$ and $Y$ have the subjects as rows and they are organized such that the rows are corresponding. For instance, the first row in $Y$ contains the behavioural information of the subject in the first row of $X$. Then, we compute the correlation matrix $R$ such that $R = XY^T$. Then, $R$ is decomposed using SVD which yields a set of uncorrelated latent variables (LVs) (***Krishnan et al., 2011***; ***Zeighami et al., 2019***). Each LVs is composed of a vector of behavioural scores, a vector of brain scores and a singular value (often referred to as saliences). The behavioural and brain scores contain the weights such that the original brain a behavioural variable maximally covary and the singular value represents the proportion of the covariance captured by this LV (***Krishnan et al., 2011***; ***Zeighami et al., 2019***).

The significance of the patterns of covariance uncovered by the LVs can be assessed using permutation testing. Permutation testing involves randomly shuffling without replacement the rows of the brain variables matrix $X$ to destroy its associations with the behavioural variables in $Y$ (***Patel et al., 2020***; ***McIntosh and Lobaugh, 2004***; ***Zeighami et al., 2019***). PLSC is applied to each shuffled brain-derived matrix with the unchanged behavioural variables as described above. The singular values obtained from every permutation form a null distribution to which the original LVs' singular values can be compared and assign a non-parametric P-value (***McIntosh et al., 1996***; ***Krishnan et al., 2011***; ***Zeighami et al., 2019***; ***McIntosh and Lobaugh, 2004***; ***Patel et al., 2020***).

The stability, reliability of specific brain scores elements or weight are assessed using bootstrap sampling (***Krishnan et al., 2011***; ***Zeighami et al., 2019***; ***McIntosh and Lobaugh, 2004***). In bootstrap sampling, the rows of both $X$ and $Y$ are randomly shuffled with replacement to create a bootstrap sample (***Efron and Tibshirani, 1986***; ***McIntosh and Lobaugh, 2004***). PLSC is then applied on every bootstrap sample to obtain a set of brain saliences as well as behaviour saliences vectors for every latent variable (***Patel et al., 2020***). We divide the brain and behavioural saliences of the original unshuffled LVs by the standard error of the bootstrap samples' saliences to obtain a bootstrap ratio (BSR). The BSR is then used to assess the significance of specific brain saliences (***Krishnan et al., 2011***; ***Zeighami et al., 2019***; ***McIntosh and Lobaugh, 2004***; ***Patel et al., 2020***).

#### Sex diferences in individual-level variation in microstructure & behaviour

As the female sample in this study has a slightly higher mean age than the males (mean female age = 29.01±3.62, mean male age = 27.71±3.67), the correlation patterns between the significant LVs with age and sex might be affected. For instance, a true correlation between an LV with age might also drive a correlation between the LV and sex or vice versa due to the previously noted bias in the sample.

To investigate further the effect of sex in our LVs, we performed the same OPNMF followed by PLS on males and females independently.

We found that for the left hemishpere, there was no significant difference between males and females in the striatum parcellation. Hence, we ran the PLS analysis for the left hemisphere without the sex as a variable as any effect of sex would not be related to microstructural variation. We obtained two significant LVs ($p<0.05$) which are shown in ***Figure 1***. We observed similar LVs as when the sex variable was included were the left LV1 mostly shows an effect of age and left LV2 shows an effect of the motor behaviours.

As the right striatum parcellation was slightly different between males and females, we conducted the PLS analysis independently between males and females. We found two significant LVs ($p<0.05$) for the female group and two significant LVs ($p<0.05$) for the male group, which can be found in ***Figure 2***. We observe that the microstructural partterns uncovered by the right LVs were different between males and females (***Figure 2***B). We hypothesize that these differences may be due to the difference in the right striatum parcellation between males and females. However, the behavioural patterns were highly similar between the two groups. Indeed, the right LV1 for males (***Figure 2***A second column) and females (***Figure 2***A second column) show mostly an effect of age and motor related behaviours while the right LV2 (***Figure 2***A third and fourth columns) shows a stronger effect of impulsivity related behaviours, similar to the previous LVs where the males and females were combined.

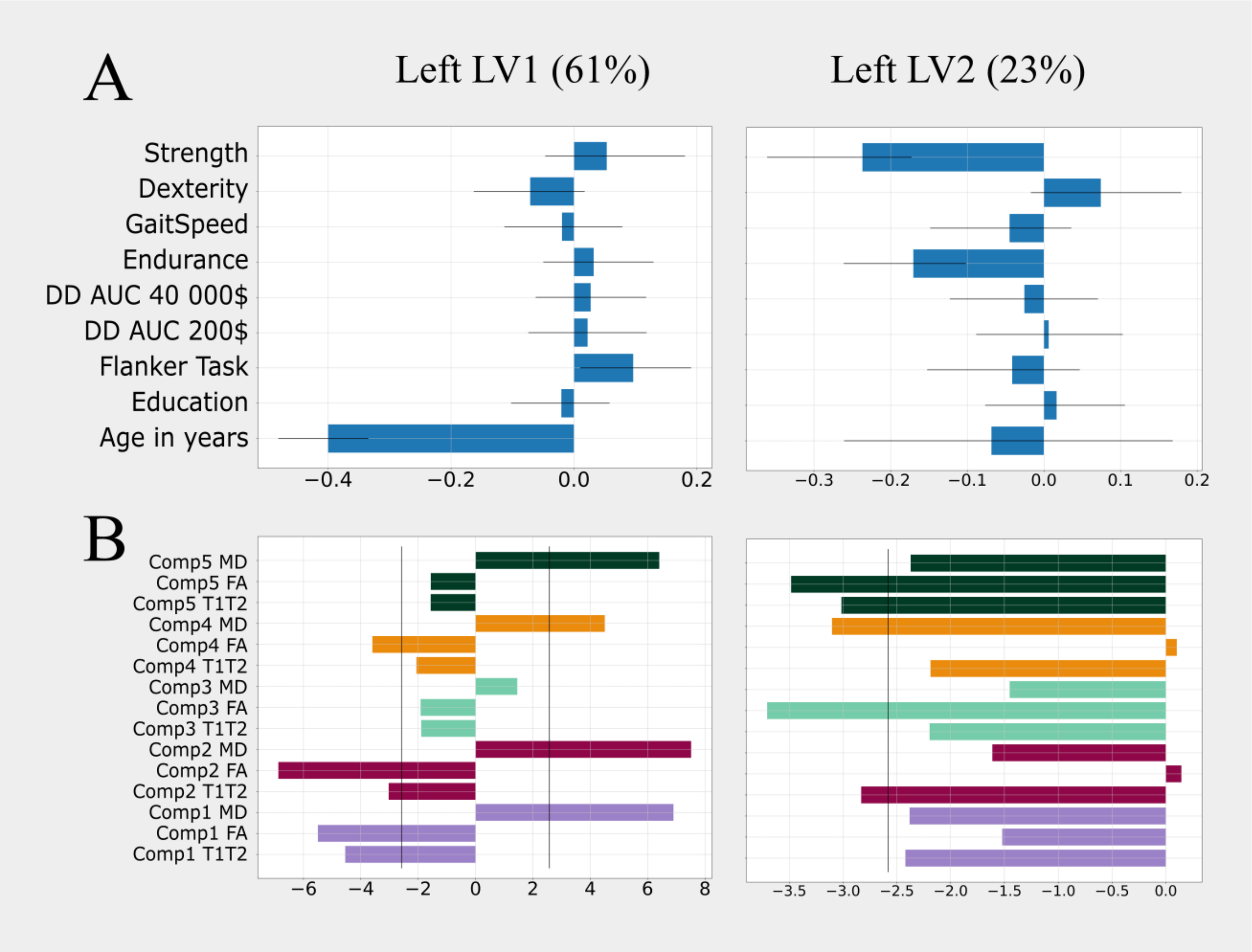


**Figure 1.** Left striatum results of the PLS analysis without the sex variable, we show only the latent variables (LVs) that were significant (p<0.05). The percentage next to the LV's name corresponds to the covariance explained by this LV. A) Behavioural patterns of the left LV1 (first column), left LV2 (second column). The y-axis denotes the behavioural and demographics measures used in the analysis (DD AUC: Delay discounting area under the curve), while the x-axis corresponds to the correlation of the behaviours with the LV. B) Microstructural patterns associated with the two significant LVs identified. Here,the y-axis correspond to the component-metric pairs and the x-axis denotes the bootstrap ratio (BSR). The black line in the microstructural patterns graph represent a BSR of 2.58 (equivalent to a 99% C.I.). The colors of the bars are associated with the component (see Figure 3C in the main manuscript).

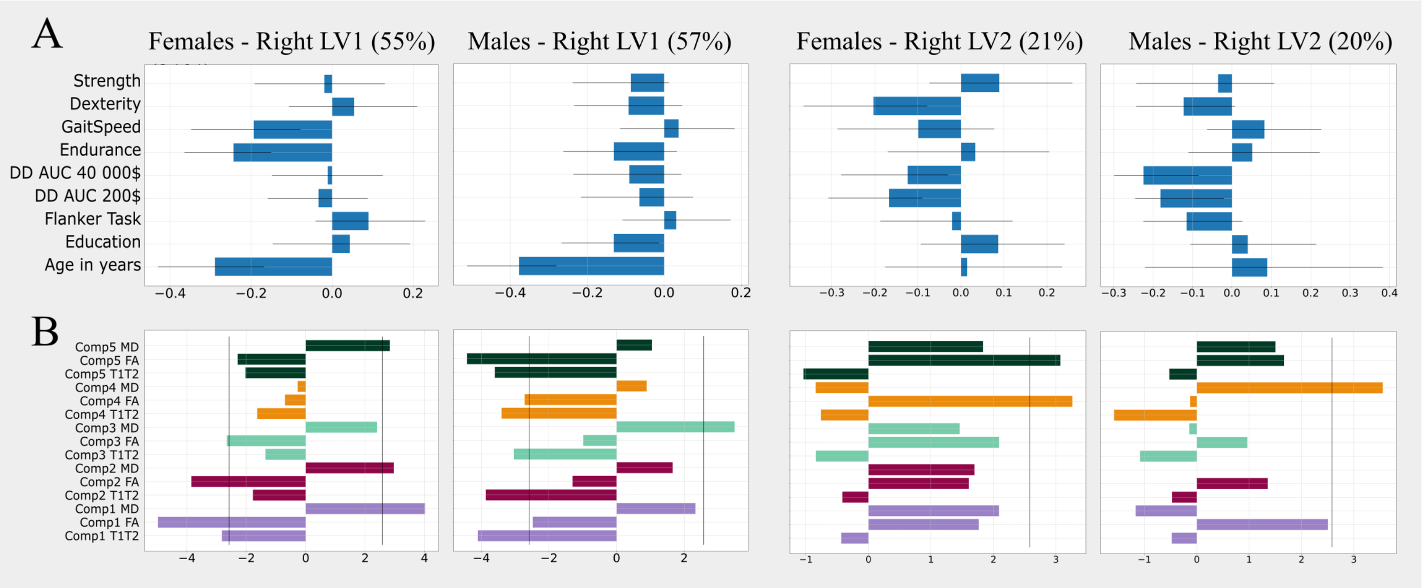


**Figure 2.** Right striatum results of the PLS analysis performed on males and females independently, we show only the latent variables (LVs) that were significant (p<0.05). The percentage next to the LV's name corresponds to the covariance explained by this LV. A) Behavioural patterns of the females right LV1 (first column), females right LV2 (third column), males right LV1 (second column) and males right LV2 (fourth column). The y-axis denotes the behavioural and demographics measures used in the analysis (DD AUC: Delay discounting area under the curve), while the x-axis corresponds to the correlation of the behaviours with the LV. B) Microstructural patterns associated with the four significant LVs identified. Here,the y-axis correspond to the component-metric pairs and the x-axis denotes the bootstrap ratio (BSR). The black line in the microstructural patterns graph represent a BSR of 2.58 (equivalent to a 99% C.I.). The colors of the bars are associated with the component (see Figure 3C in the main manuscript).

### References

**Boutsidis C**, Gallopoulos E. SVD based initialization: A head start for nonnegative matrix factorization. Pattern Recognition. 2008; 41(4):1350–1362. http://www.sciencedirect.com/science/article/pii/S0031320307004359, doi: https://doi.org/10.1016/j.patcog.2007.09.010.

**Efron B**, Tibshirani R. Bootstrap methods for standard errors, confidence intervals, and other measures of statistical accuracy. Statistical science. 1986; p. 54–75.

**Krishnan A**, Williams LJ, McIntosh AR, Abdi H. Partial Least Squares (PLS) methods for neuroimaging: A tutorial and review. NeuroImage. 2011; 56(2):455 – 475. http://www.sciencedirect.com/science/article/pii/S1053811910010074, doi: https://doi.org/10.1016/j.neuroimage.2010.07.034, multivariate Decoding and Brain Reading.

**McIntosh AR**, Lobaugh NJ. Partial least squares analysis of neuroimaging data: applications and advances. NeuroImage. 2004; 23:S250 – S263. http://www.sciencedirect.com/science/article/pii/S1053811904003866, doi: https://doi.org/10.1016/j.neuroimage.2004.07.020, mathematics in Brain Imaging.

**McIntosh A**, Bookstein F, Haxby JV, Grady C. Spatial pattern analysis of functional brain images using partial least squares. Neuroimage. 1996; 3(3):143–157.

**Patel R**, Steele CJ, Chen AGX, Patel S, Devenyi GA, Germann J, Tardif CL, Chakravarty MM. Investigating microstructural variation in the human hippocampus using non-negative matrix factorization. NeuroImage. 2020; 207:116348. http://www.sciencedirect.com/science/article/pii/S1053811919309395, doi: https://doi.org/10.1016/j.neuroimage.2019.116348.

**Sotiras A**, Resnick SM, Davatzikos C. Finding imaging patterns of structural covariance via Non-Negative Matrix Factorization. NeuroImage. 2015; 108:1–16. https://pubmed.ncbi.nlm.nih.gov/25497684https://www.ncbi.nlm.nih.gov/pmc/articles/PMC4357179/https://www.ncbi.nlm.nih.gov/pmc/articles/PMC4357179/pdf/nihms649302.pdf, doi: 10.1016/j.neuroimage.2014.11.045.

**Varikuti DP**, Genon S, Sotiras A, Schwender H, Hoffstaedter F, Patil KR, Jockwitz C, Caspers S, Moebus S, Amunts K. Evaluation of non-negative matrix factorization of grey matter in age prediction. Neuroimage. 2018; 173:394–410. https://www.ncbi.nlm.nih.gov/pmc/articles/PMC5911196/pdf/nihms952495.pdf.

**Zeighami Y**, Fereshtehnejad SM, Dadar M, Collins DL, Postuma RB, Mišić B, Dagher A. A clinical-anatomical signature of Parkinson's disease identified with partial least squares and magnetic resonance imaging. NeuroImage. 2019; 190:69 – 78. http://www.sciencedirect.com/science/article/pii/S1053811917310741, doi: https://doi.org/10.1016/j.neuroimage.2017.12.050, mapping diseased brains.
